# Characterization of Distinct Monocyte Subtypes and Immune Features Associated with HIV, Tuberculosis, and Coronary Artery Disease in a Ugandan Cohort Using Mass Cytometry

**DOI:** 10.1101/2025.08.25.672210

**Authors:** José Cobeña-Reyes, Celestine N. Wanjalla, Manuel G. Feria, Joshua Simmons, Tecla Temu, Cindy Nochowicz, Sheikh Yasir Arafat, Cissy Kityo, Geofrey Erem, Christopher T. Longenecker, Sandra Andorf, Moises A. Huaman

**Affiliations:** Division of Biomedical Informatics, Cincinnati; Children’s Hospital Medical Center, Cincinnati, OH, USA; Division of Infectious Diseases, Vanderbilt University Medical Center, Nashville, TN, USA; The Center for AIDS Health Disparities Research, Nashville, TN, USA; Veteran’s Health Administration, Tennessee Valley Healthcare System, Nashville, TN 37212, USA; Division of Infectious Diseases, University of Cincinnati College of Medicine, Cincinnati, OH, USA; Department of Pathology, Mass General Brigham, Harvard Medical School, Boston, MA, USA; Department of Biostatistics, Health Informatics, and Data Sciences, College of Medicine, University of Cincinnati, Cincinnati, OH, USA; Joint Clinical Research Centre, Kampala, Uganda; Makerere University, Kampala, Uganda; Division of Cardiology, Department of Global Health, University of Washington, Seattle, WA, USA; Division of Allergy and Immunology, Cincinnati Children’s Hospital Medical Center, Cincinnati, OH, USA; Department of Pediatrics, University of Cincinnati College of Medicine, Cincinnati, OH, USA

## Abstract

Coronary artery disease (CAD), tuberculosis (TB), and HIV represent major global health burdens. Individuals affected by one or more of these conditions often exhibit chronic inflammation and immune dysregulation, with monocytes playing a central role in these processes. Monocyte subsets are known to expand in individuals with HIV, TB, or CAD. However, the precise mechanisms by which these cells contribute to inflammation and immune responses in the context of these conditions remain poorly understood. In this study, we employed high-dimensional mass cytometry to characterize monocyte heterogeneity in 61 Ugandan adults with varying combinations of HIV, latent TB, and subclinical or overt CAD. Through an integrative approach combining manual gating, unsupervised clustering, and elastic net penalization, we identified distinct monocyte phenotypes associated with CAD and TB. Importantly, individuals with CAD, especially those with more extensive disease (Segment Involvement Score >2), showed reduced surface expression of the anti-inflammatory scavenger receptor CD163 on non-classical monocytes. Notably, unsupervised clustering further revealed two distinct non-classical monocyte subsets associated with disease states: A CD86^dim^ CX3CR1^dim^ CD45RA^+^ GPR56^+^ CXCR3^+^ subset significantly depleted in individuals with CAD, and a CD86^+^ CX3CR1^++^ CD45RA^++^ GPR56^−^ CD38^−^ CXCR3^−^ subset enriched in individuals with TB. These findings underscore the complexity of the monocyte landscape in CAD progression, particularly within settings of HIV and TB co-endemicity. We hope this work motivates further research and offers insights for the development of new precision biomarkers and immune-targeted therapies to prevent or treat CAD, TB, and HIV in populations.

## INTRODUCTION

HIV,^1^ tuberculosis (TB),^2^ and cardiovascular diseases^3,4^ are major public health concerns and leading causes of morbidity and mortality worldwide. Individuals residing in Sub-Saharan Africa are particularly vulnerable^5–7^ , often facing the added challenge of living with one or more of these comorbid conditions simultaneously. People living with HIV (PWH) experience complex immune dysregulation despite appropriate antiretroviral therapy, which may contribute to various comorbidities such as coronary artery disease (CAD)^8^ or increased susceptibility to other infectious diseases such as TB.^9^ Similarly, people with TB exhibit altered immune responses that can exacerbate immune dysfunction and further elevate CAD risk.^10,11^

Monocytes are cells of the innate immune system that play a critical role in the response to infection and in regulating the immune response. Monocytes are classified based on their CD14 and CD16 expression as Classical Monocytes (CD14++ CD16-), Intermediate monocytes (CD14++ CD16+) and non-Classical monocytes (CD14+ CD16++). In people living with HIV, TB, or CAD, monocyte populations undergo significant alterations in both frequency and function.^12–15^ In CAD, lower levels of classical monocytes circulating in the blood are usually encountered.^16^ Intermediate monocytes tend to aggregate and form monocyte-platelet aggregates in people with ST-elevation myocardial infarction due to coronary atherothrombosis.^17^ Further, monocytes infiltrate plaque and differentiate into macrophages (i.e. monocyte-derived macrophages). However, these macrophages exhibit a decreased ability to migrate, promoting atherogenesis.^18,19^ In TB infection, CD16+ monocyte subsets are expanded, reflecting alterations in the composition of circulating monocyte populations.^20^ The expansion is, however, reversed by anti-TB treatment.^20^ Classical monocytes that express inflammatory markers typically increase as the disease progresses, whereas the percentage of non-classical monocytes expressing anti-pathogen infection markers decreases.^21^ In PWH, alterations in the distribution of monocyte subsets are also observed, including an increase in intermediate and non-classical monocytes.^22^ Interestingly, intermediate monocyte subsets are preferentially infected with HIV due to the facilitation of viral entry.^23,24^ Moreover, there are indications that CD16+ subsets might serve as a reservoir for HIV, complicating viral suppression.^25^ Understanding changes in the distribution and phenotype of monocyte subsets is crucial for uncovering disease mechanisms and potential therapeutic targets.

In this work, to characterize monocyte populations in individuals with HIV, TB, and CAD, a cohort of people living with these conditions was studied. Using mass cytometry data, both manually gated monocyte subtypes and those identified through unsupervised clustering were examined, and their distributions were compared across groups defined by TB, HIV, and CAD status. Additionally, elastic net regression was employed to identify monocyte features most strongly associated with CAD status. This statistical approach enabled the identification of previously undefined immune features that may serve as correlates for disease progression and potential therapeutic intervention.

Given the critical role of monocyte subsets in the regulation of HIV, TB, and CAD, this study aimed to further uncover critical insights into how monocyte populations are altered in HIV, TB and CAD, by leveraging high-parameter mass cytometry data and advanced statistical analysis.

## METHODS

### Study design and participants

Mass cytometry data from manually-gated monocytes isolated from peripheral blood mononuclear cells (PBMCs) of 61 participants were analyzed. All the participants were enrolled at the Joint Clinical Research Center in Kampala, Uganda, in the Ugandan study of HIV effects on the myocardium and atherosclerosis (mUTIMA). For this analysis, the deidentified dataset contained information on individuals living with or without HIV, TB, and CAD. The protocol of the parent study was previously described.^26^ The study was approved by the University Hospitals Cleveland Medical Center Institutional Review Board, the Joint Clinical Research Centre Research Ethics Committee, and the Uganda National Council for Science and Technology. All participants signed written informed consent.

TB infection status was defined as latent tuberculosis infection (TBI+) or negative (TB-) based on comprehensive TB symptom questionnaires and interferon-gamma release assay (IGRA) testing (QuantiFERON®-TB). Previous active tuberculosis (TBpr) was defined by self-reported history of prior active TB and completion of TB treatment, which was confirmed by review of medical records. HIV+/-was defined based on FDA-approved HIV testing. The Atherosclerotic Cardiovascular Disease (ASCVD) pooled cohort equations (PCE) were used to calculate 10-year ASCVD risk scores.^27^ All participants underwent coronary computed tomography angiography to quantify the presence and severity of CAD.

Two types of categorizations were employed based on the CAD status: A binary classification defined as the presence or absence of coronary plaque (CAD-/+) and a categorization based on burden of disease using the segment involvement score (SIS), an integer-based measure that assigns a value of 1 to each coronary artery segment with detectable atherosclerotic plaque, irrespective of plaque size or degree of luminal obstruction^28^: CAD-, CAD+ minimum (CAD min): ≤ 2 SIS; and CAD+ greater than minimum (CAD+ >min): > 2 SIS. Five study groups were also defined by combining the TB and HIV status as HIV-TB-, HIV-TBI+, HIV+TB-, HIV+TBI+, and HIV+TBpr.

### Mass cytometry

Cryopreserved PBMCs were rapidly thawed in a 37 °C water bath and immediately transferred to 10 mL ice-cold RPMI-1640 (no serum) containing 20 μL Nuclease S7 (Roche) to degrade extracellular nucleic acids. Cells were washed twice by centrifugation at 420 × g for 10 min at room temperature (RT) in 10 mL phosphate-buffered saline (PBS; without Ca²⁺/Mg²⁺), decanted, and gently resuspended in the residual supernatant. Viable cells were counted by trypan-blue exclusion on a hemocytometer, and aliquots of 3 × 10⁶ cells were transferred to 5 mL polystyrene tubes (Falcon) for staining.

Cells were washed once in 2 mL PBS/1% bovine serum albumin (BSA; Sigma-Aldrich) at 420 × g × 10 min (RT) and then stained with 2 μL Live/Dead Rhodium-103 (Fluidigm; 1:500 final) for 15 min at 37 °C. After two additional washes in PBS/1% BSA (as above), pellets were tapped dry, then resuspended in 50 μL staining buffer (PBS/1% BSA). A master mix of metal-conjugated surface antibodies (see Table S1) and EQ™ Four Element Calibration Beads (Fluidigm) was prepared to a volume of 40 μL per sample, brought to ∼100 μL total staining volume with staining buffer, and added to each tube. Samples were incubated for 30 min at RT, washed twice in 2 mL PBS (420 × g × 10 min, RT), and fixed by adding 20 μL 16% paraformaldehyde (PFA; Thermo Fisher) to 200 μL cells (final 1.6% PFA), incubating 15 min at RT.

Fixed cells were washed once in 2 mL PBS (800 × g × 7 min, RT), aspirated, and permeabilized by adding 1 mL of ice-cold methanol (MeOH), then stored at –20 °C overnight. Cells were then labeled with 2 μL of 25 μM iridium DNA intercalator (Fluidigm) in the presence of 1.6% PFA (20 μL PFA added to 200 μL cells) for 20 min at RT, followed by an overnight incubation at 4 °C.

Immediately before acquisition, cells were washed once in PBS and once in Millipore-grade H₂O (800 × g × 7 min), resuspended at 5 × 10⁵ cells/mL in double-distilled H₂O, and spiked with equilibration (EQ) beads at a 1:10 bead-to-cell ratio (vortex beads vigorously before use). Finally, cell suspensions were passed through a 35 μm nylon filter (CellTrics, Sysmex) and acquired on the Helios at <500 events.

### Monocyte phenotyping: Manual gating

Conventional monocyte subsets were manually gated from PBMCs into total (TM), classical (CM), intermediate (IM), and non-classical (nCM) categories based on CD14 and CD16 expression, as previously described^6^ (see Supplemental Figure S1).

For each of the monocyte subsets, several populations were identified by manual gating. For the TM, frequencies of the following populations were determined: CD14+, CD14+ HLA-DR+, CD14+ CX3CR1+, CD14+ CD86+, CD14+ CD163+. For the CM, the following subpopulations were considered: CD14+ CD16-, CD14+ CD16- CX3CR1+, CD14+ CD16- CD86+, CD14+ CD16- CD163+. Subpopulations based on the same markers were studied for IM (CD14+ CD16-, CD14+ CD16- CX3CR1, CD14+ CD16- CD86+, CD14+ CD16- CD163+) and nCM (CD14- CD16+, CD14- CD16+ CX3CR1+, CD14- CD16+ CD86+, CD14- CD16+ CD163+).

Additionally, the Median Signal Intensity (MSI) of markers HLA-DR, CX3CR1, and CD86 was computed for TM, CM, IM, and nCM. MSI of CD163 was computed for TM, IM, and nCM.

### Monocyte phenotyping: Unsupervised clustering

FCS files containing the total monocytes (TM) obtained from manual gating were read using the flowCore^29^ package from the Bioconductor^30^ project, and transformed using the *arcsinh* function with a cofactor of 5.^31^ Ten thousand cells were sampled randomly from each file. If a file contained fewer than 10,000 cells, all cells were used. Of the files, 46 contained more than 10,000 cells, while 15 contained fewer. Unsupervised clustering via FlowSOM^32^ with default parameters and a predetermined number of 12 clusters was used to identify phenotypically distinct monocyte populations. All markers were used for unsupervised clustering. For each of the samples, the percentage of cells assigned to each of these clusters (i.e. monocyte populations) was calculated.

To determine the marker expression within each monocyte population, scaled median expression values^33^ were visualized using a heatmap. To visualize 2D map projections of the cell populations, the stratified CAD/SIS study groups (CAD-, CAD+ min and CAD+ > min), were used. For each group, 80,000 cells were randomly sampled and plotted using Uniform Manifold Approximation Projections (UMAPs) with colors for the populations based on the unsupervised clustering overlayed.^34^

### Elastic net regularization

The features from manual gating and the population percentages from FlowSOM were used to fit a generalized linear model, incorporating a regularization step through elastic net^35^, to identify the features most strongly associated with CAD status. In this model, due to the small sample size, the CAD-/+ classification was used as the predicted variable. In the context of CAD status, negative coefficients from the elastic net model are primarily associated with CAD– (absence of disease), while positive coefficients are associated with CAD+ (presence of disease). Elastic net regularization combines the penalties of Ridge and LASSO regression, enabling both variable selection and shrinkage, which helps reduce overfitting and isolate the most informative features. Alpha values ranging from 0 (Ridge Regression) to 1 (LASSO regression), in increments of 0.1 were tested. For each value of alpha, one hundred values of lambda were tested using a 10-fold cross-validation to select the optimal lambda based on the smallest mean misclassification error (MME). Coefficients of non-zero features, i.e. variables retained in the regularized model, were visualized in a bar plot.

### Statistical analysis

Percentages of populations and MSI of the markers from the manual gating, and the percentage of populations obtained from unsupervised clustering were compared across the study groups using a Kruskal-Wallis test.^36^ The p-values were adjusted to control the false discovery rate (FDR) using the Benjamini-Hochberg method^37^. For clusters with a significant Kruskal-Wallis test (FDR-adjusted p < 0.05), Wilcoxon tests^38^ between each pair of study groups were performed.

All statistical analyses were performed in R (version 4.4.2). Boxplots show the medians, the 1st and 3rd quartiles. Whiskers represent the minimal and maximal values after excluding outliers, defined as points beyond 1.5 times the interquartile range (IQR) from the 25th or 75th percentile. Individual data points representing each participant are shown. Plotting unless otherwise stated was done using the ggplot2 R package (version 3.5.1). R packages flowCore (version 2.18.0), FlowSOM (version 2.14.0), glmnet (version 4.1-8), ggpubr (version 0.6.0), umap (version 0.2.10.0), corrplot (version 0.95), and ComplexHeatmap (version 2.22.0) were also employed.

## RESULTS

### Study population

A total of 61 participants were grouped into CAD-/+ classification: CAD- (n=40), CAD+ (n=21), (Table S2); and into the stratified CAD/SIS classification: CAD– (n = 40), CAD+ min (n = 13), and CAD+ >min (n = 8) (Table 1). The demographics and clinical characteristics of these participants categorized by HIV and TB status are shown in Table 2.

**Table 1:** Demographics and clinical characteristics of the cohort using 3 CAD/SIS groups: CAD-, CAD+ min, and CAD+ >min.

|  | *All patients*<br>*N = 61* | *CAD-*<br>*N = 40,*<br>*(65.6%)* | *CAD+ min*<br>*N = 13,*<br>*(21.3%)* | *CAD+ >min*<br>*N = 8,*<br>*(13.1%)* | *p-value* |
| --- | --- | --- | --- | --- | --- |
| **Age [years]** | 61 (56, 65) | 60 (55, 65) | 60 (57, 64) | 65 (64, 72) | 0.068 |
| **Sex: female, n (%)** | 23 (37.7) | 17 (42.5) | 5 (38.5) | 1 (12.5) | 0.38 |
| **ASCVD risk** | 9.4 (5.7, 15.2) | 10.6 (5.3, 14.3) | 8.0 (5.0, 12.7) | 11.0 (7.5, 27.1) | **0.03** |
| **BMI [pound/in<sup>2</sup>]** | 29.2 (25.6, 33.3) | 29.0 (26.2, 32.8) | 30.5 (25.5, 34.2) | 30.1 (26.6, 35.1) | 0.81 |
| **Weight [pounds]** | 176.4 (151.0, 196.2) | 176.4 (156.5, 196.2) | 176.4 (149.9, 194.0) | 174.2 (157.6, 191.8) | 0.87 |
| **Height [in]** | 64.4 (61.8, 66.3) | 64.6 (62.0, 67.3) | 64.0 (61.0, 64.8) | 64.0 (62.7, 65.4) | 0.28 |
| **Continuous variables are presented as median (interquartile range)** |  |  |  |  |  |

**Table 2:** Demographics and clinical characteristics of the cohort classified using five HIV/TB groups: HIV-TB-, HIV-TBI+, HIV+TB-, HIV+TBI+, HIV+TBpr.

|  | *HIV- TB-*<br>*N = 13*<br>*(21.3%)* | *HIV- TBI+*<br>*N = 22*<br>*(36.1%)* | *HIV+ TB-*<br>*N = 9*<br>*(14.8%)* | *HIV+ TBI+*<br>*N = 9*<br>*(14.8%)* | *HIV+ TBpr*<br>*N = 8*<br>*(13.0%)* | *p-*<br>*value* |
| --- | --- | --- | --- | --- | --- | --- |
| **Age [years],** | 60.0 (56.5, 63.8) | 62.0 (54.5, 63.8) | 65.0 (60.0, 66.0) | 56.5 (54.3, 59.0) | 62.5 (60.3, 65.0) | 0.25 |
| **Sex: female, n (%)** | 8 (66.7) | 7 (31.8) | 2 (22.2) | 3 (33.4) | 3 (37.6) | 0.26 |
| **ASCVD risk** | 15.3 (5.4, 24.8) | 9.7 (4.9, 12.7) | 10.6 (7.8, 12.1) | 7.5 (4.3, 9.8) | 9.8 (7.5, 33.7) | 0.38 |
| **BMI [pound/in<sup>2</sup>]** | 28.2 (26.9, 32.0) | 31.8 (27.6, 34.2) | 26.1 (25.3, 29.2) | 28.1 (25.6, 34.0) | 29.6 (27.5, 34.1) | 0.55 |
| **Weight [pounds]** | 177.5, (154.9, 185.7) | 174.2 (159.3, 198.4) | 152.1 (129.0, 185.2) | 174.2 (157.1, 187.4) | 189.6 (167.8, 199.0) | 0.59 |
| **Height [in]** | 65.4 (62.4, 68.2) | 63.8 (61.8, 65.1) | 62.2 (60.2, 65.4) | 66.1 (63.6, 66.7) | 65.2 (62.9, 66.5) | 0.48 |
| **Continuous variables are presented as median (interquartile range)** |  |  |  |  |  |  |

### CAD+ >min participants show lower MSI of CD163 than CAD- and CAD+ min in manually gated nCM

Frequencies of different subpopulations and MSI of several markers in TM, CM, IM, and nCM, based on manual gating, were compared across the CAD-/+ and CAD/SIS groups (Supplementary Figures S2-S9). Statistically significant differences were observed in the nCM (Supplementary Figure S5, FDR-adjusted p = 0.020). Individuals in the CAD+ >min group had a lower MSI of CD163 compared to both the CAD- and CAD+ min groups (p = 0.023 in both cases). No statistically significant differences in the manually gated populations were observed across the five TB/HIV nor in the CAD-/+ groups (Supplementary Figures S6 – S13).

### Unsupervised clustering identifies distinct subpopulations

Unsupervised clustering via FlowSOM was used to separate the total monocytes into 12 clusters. Scaled median marker expressions were used to classify these into 2 CM, 5 IM, and 5 nCM populations (Supplementary Figure S14). The 2 CM populations (clusters 1 and 3) were merged into one population as relevant markers such as CD14, CD16, and HLA-DR were similarly expressed. Four IM subpopulations (clusters 4 and 5; and clusters 2 and 9) were also combined into two populations. Three nCM populations (clusters 6, 7, and 10) were merged due to similar characteristics. This resulted in a final list of 7 clusters, corresponding to 1 CM, 3 IM, and 3 nCM populations, for downstream analyses (Figure 1). The heatmap of scaled marker expression of each population, along with their percentages in all cells, is shown in Figure 1A. Visualization of the clustering results in a UMAP (Figure 1B) showed that most populations clustered closely together. However, two of the nCM populations were slightly (purple population in the plot) or markedly (orange population) separated from the rest.

**Figure 1:**
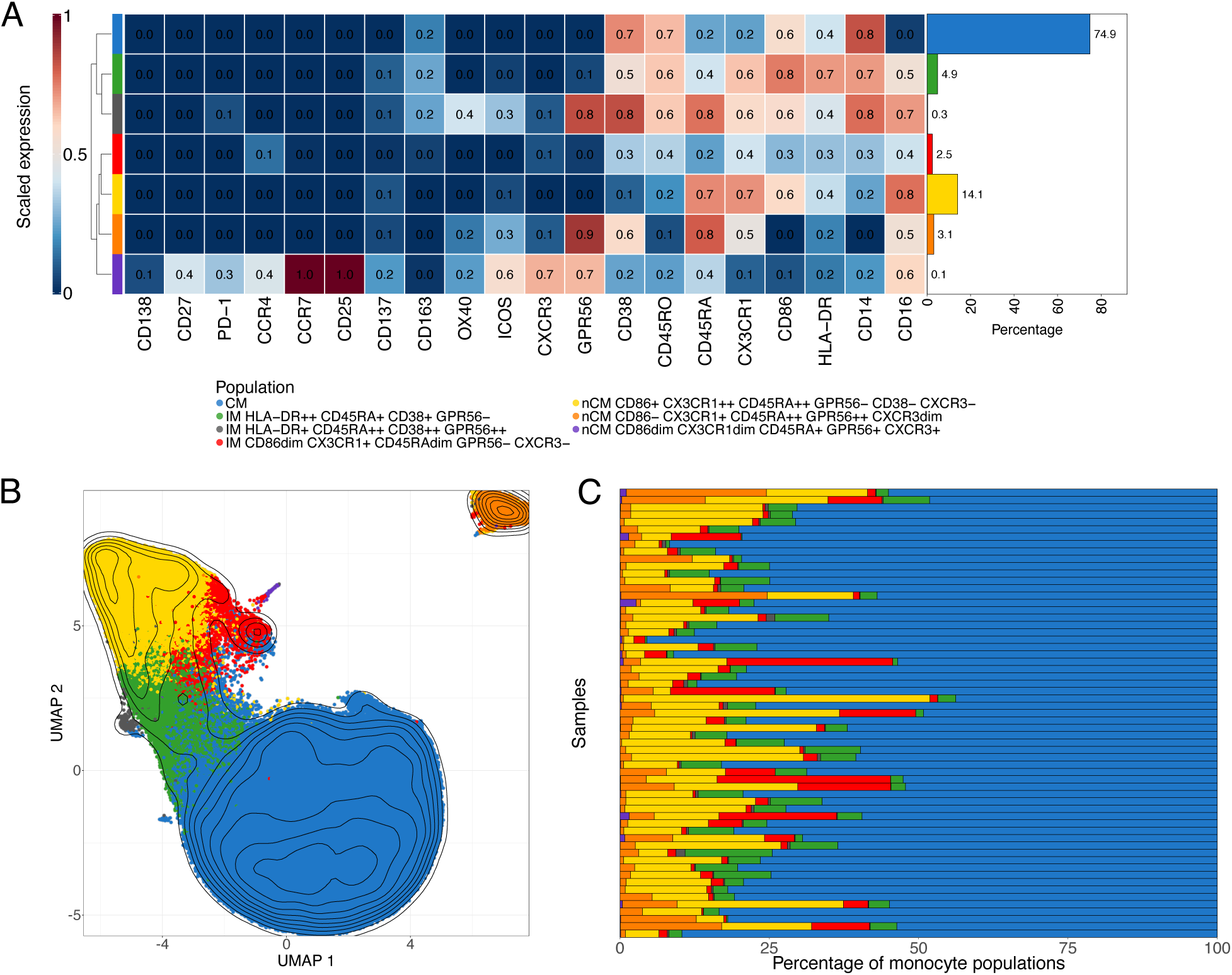
Monocyte subpopulations identification via FlowSOM and UMAP. **(A)** Scaled median marker expressions of the monocyte populations found using FlowSOM. Seven populations (1 classical, 3 intermediate, and 3 non-classical monocytes) were identified after merging similar populations (see Supplemental Figure 13). The bar plot on the right indicates the percentage of the population with respect to all cells, i.e. all manually gated monocytes used during clustering, in all the samples. **(B)** Uniform Manifold Approximation and Projections (UMAPs) of 240000 monocytes randomly selected from the cells used in the unsupervised clustering. The population obtained from the FlowSOM analysis were overlaid. **(C)** Percentage of the monocyte populations obtained from FlowSOM in each of the samples in the cohort.

The CM population comprises the highest percentage of monocytes (74.9%), which is also visually observed in most samples (Figure 1C). Among the three IM populations, the HLA- DR++ CD45RA+ CD38+ GPR56- is the largest (4.9%), followed by the CD86dim CX3CR1+ CD45RAdim GPR56- CXCR3- (2.5%) and by HLA-DR+ CD45RA++ CD38++ GPR56++ (0.3%). The nCM portion includes the CD86+ CX3CR1++ CD45RA++ GPR56- CD38- CXCR3+ population (14.1%), which is the largest after the CM. The other two nCM populations are CD86- CX3CR1+ CD45RA++ GPR56++ CXCR3dim (3.1%), which is located separated from the other populations in the corner of the plot, possibly due to the lack of expression in CD14 and CD86; and the population CD86dim CX3CR1dim CD45RA+ GPR56+ CXCR3+ (0.1%).

### Percentages of two di_erent nCM populations from the unsupervised clustering di_er between the di_erent CAD/SIS and TB/HIV study groups

Next, the frequencies of the seven populations derived from unsupervised clustering were compared, calculated as percentages of the total monocyte population used for clustering, across our study groups.

Hypothesis testing was conducted to identify differences across the CAD/SIS groups (CAD- , CAD+ min, and CAD+ >min groups). The CD86dim CX3CR1dim CD45RA+ GPR56+ CXCR3+ nCM cluster (purple population in the UMAPs, Figure 2A) showed a statistically significant difference in the percentage of cells among the three study groups (FDR-adjusted p = 0.033). In post hoc analyses, participants from the CAD+ >min group had lower proportions of this population compared to the CAD- (p = 0.033) and CAD+ min groups (p = 0.022, Figure 2B). CAD- and CAD+ min participants showed comparable proportions of this population among total monocytes. There were no significant differences across the CAD-/+ groups (Supplementary Figure S15).

**Figure 2:**
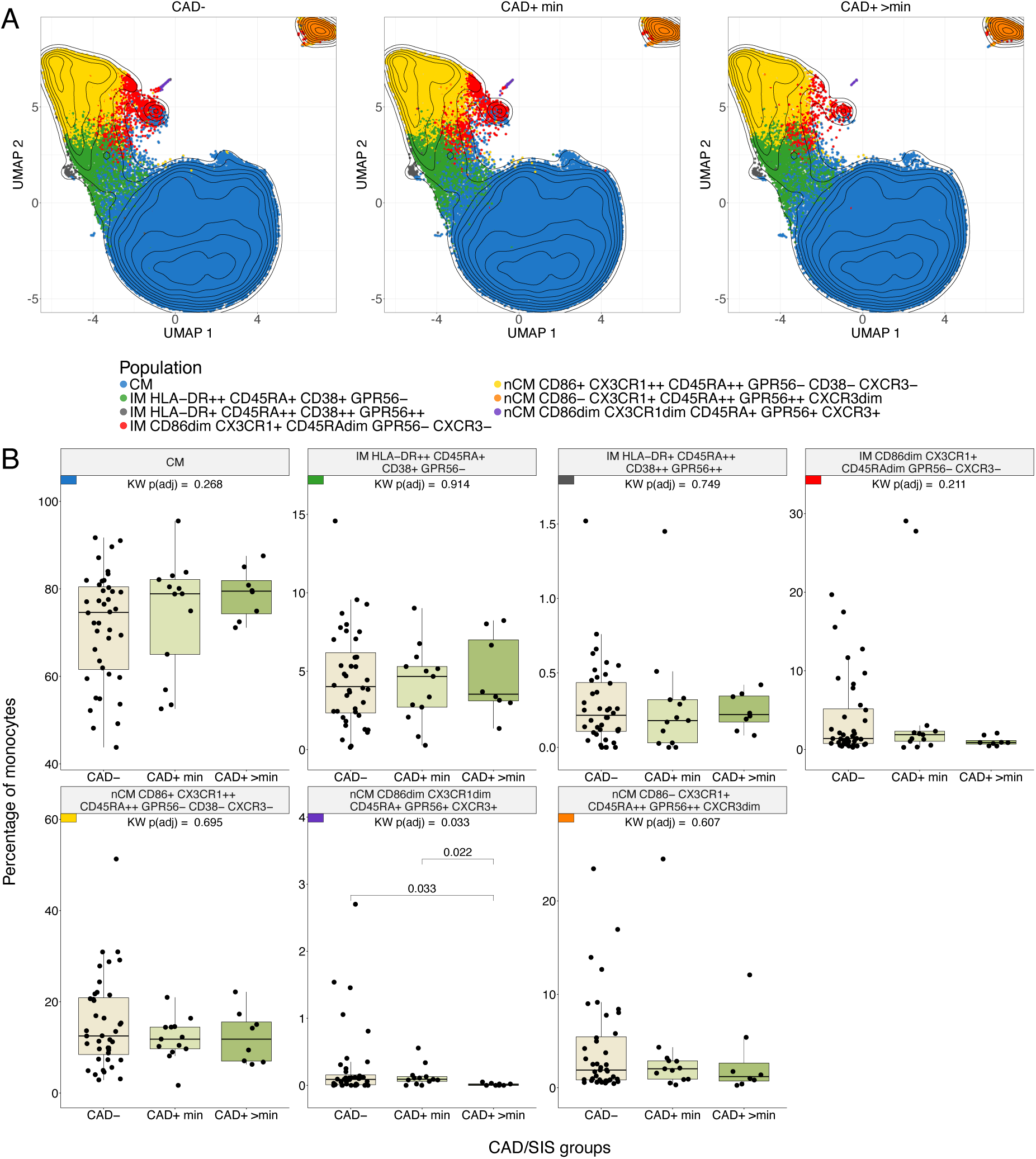
Differences in monocyte subpopulation obtained with FlowSOM across CAD/SIS Groups. **(A)** Uniform Manifold Approximation and Projections (UMAPs) stratified by CAD/SIS study groups. 80000 cells were randomly selected per CAD group, and clusters from the FlowSOM analysis were overlaid. Contour lines are of all cells sampled.**(B)** Statistical differences of percentages of cells for each monocyte population obtained from unsupervised clustering by FlowSOM across the three CAD/SIS study groups. Kruskal-Wallis (KW) tests were performed as an omnibus test to assess overall differences across the groups and p-values were adjusted for False Discovery Rate (FDR) using the Benjamini-Hochberg method. Only the population nCM CD86dim CX3CR1dim CD45RA+ GPR56+ CXCR3+ was statistically significant (FDR-adjusted p = 0.033) and was further analyzed by post-hoc Wilcoxon tests across the three groups. The percentage in the CAD+ >min study group was statistically lower than the CAD+ min (p = 0.022) and CAD- (p = 0.033).

Then, the proportion of each of the seven FlowSOM-derived populations was compared across the five HIV/TB study groups. In this analysis, a different nCM population showed statistically significant differences between the groups. The nCM cluster characterized as CD86+ CX3CR1++ CD45RA++ GPR56- CXCR3- (yellow population in the UMAPs, Figures 1 and S16) was present at significantly higher proportions in the HIV-TBI+ group compared to all the other four TB/HIV clinical groups (Figure 3). Interestingly, the same trend was observed in the manual gating results (Figure S13), where the HIV-TBI+ group showed the highest median of both percentage of CX3CR1+ and MSI of CX3CR1 in nCM. The trends are a sign of inflammation in HIV/TB co-infection,^39^ in agreement with previous research.^20^

**Figure 3:**
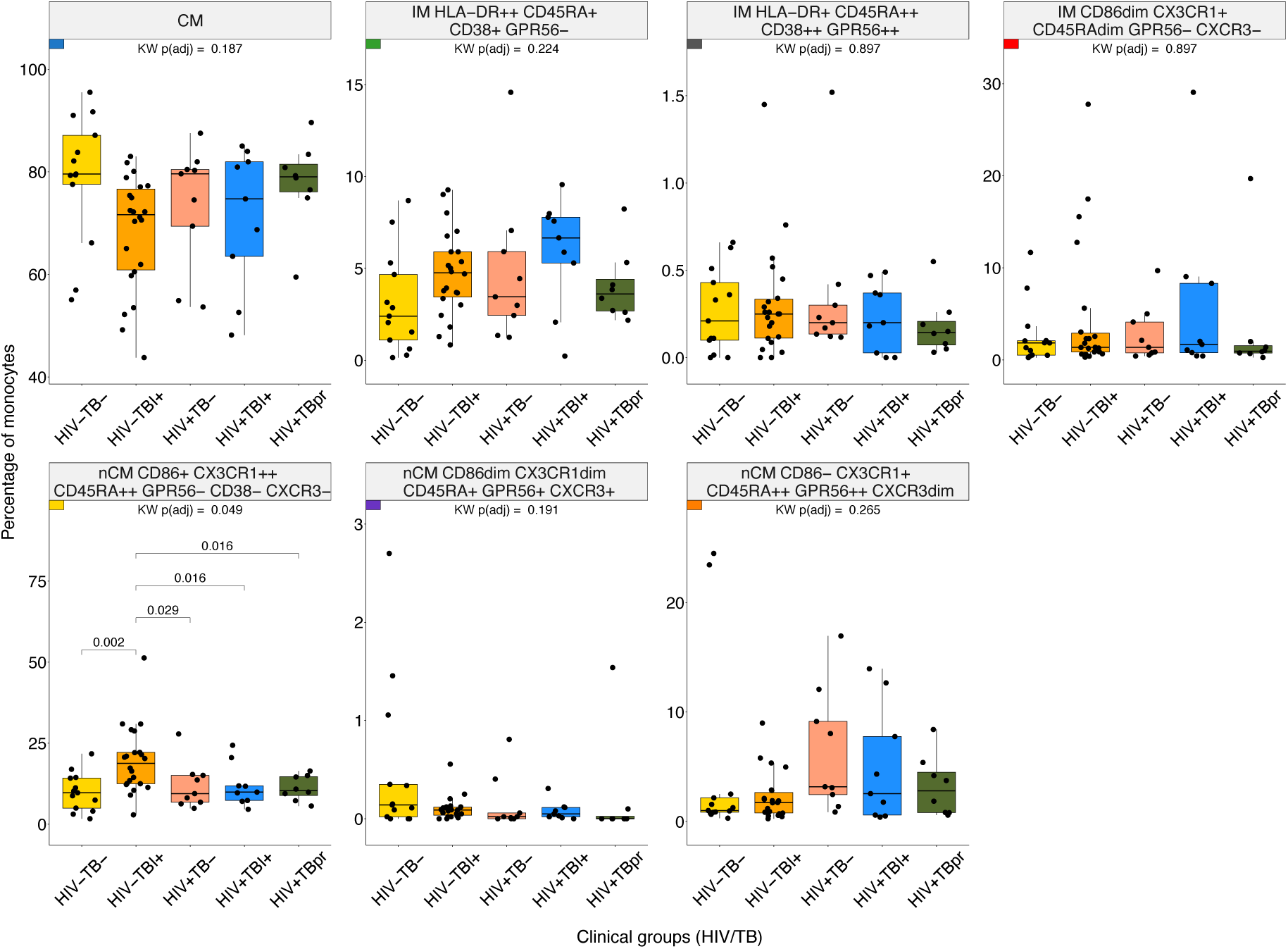
Differences in monocyte subpopulations obtained with FlowSOM across HIV/TB Groups. Statistical differences of percentages of cells for each monocyte population obtained from unsupervised clustering by FlowSOM across the five HIV/TB study groups. Kruskal-Wallis (KW) tests were performed as an omnibus test and p-values were adjusted for False Discovery Rate (FDR) using Benjamini-Hochberg method. Among the populations tested, only nCM CD86+ CX3CR1++ CD45RA++ GPR56-CD38-CXCR3-reached statistical significance (FDR-adjusted p < 0.05) and was further subjected to post-hoc Wilcoxon tests across the five groups. The small colored rectangles on top left corners follow the colors used in the UMAPs in Figures 1 and 2.

### Non-classical monocyte CX3CR1 expression and subset composition correlate with cardiovascular risk

To identify monocyte characteristics associated with CAD, elastic net regression analysis was performed using CAD-/+ status as the predicted variable. The regression model included 32 features (MSI and percent of populations) obtained by manually gating, and the percentages of the 7 populations identified in the FlowSOM clustering.

During model tuning, an alpha value of 0.6 was selected based on evaluation across 100 lambda values per alpha, as this combination yielded the lowest mean misclassification error (Figure S17). As a result, the fitted model was equivalent to an elastic net penalization that combines L1 and L2 regularization, shrinking coefficients of less important features to zero and effectively selecting a sparse set of predictors. Figure 4 shows the coefficients of the non-zero features retained in the final model.

**Figure 4:**
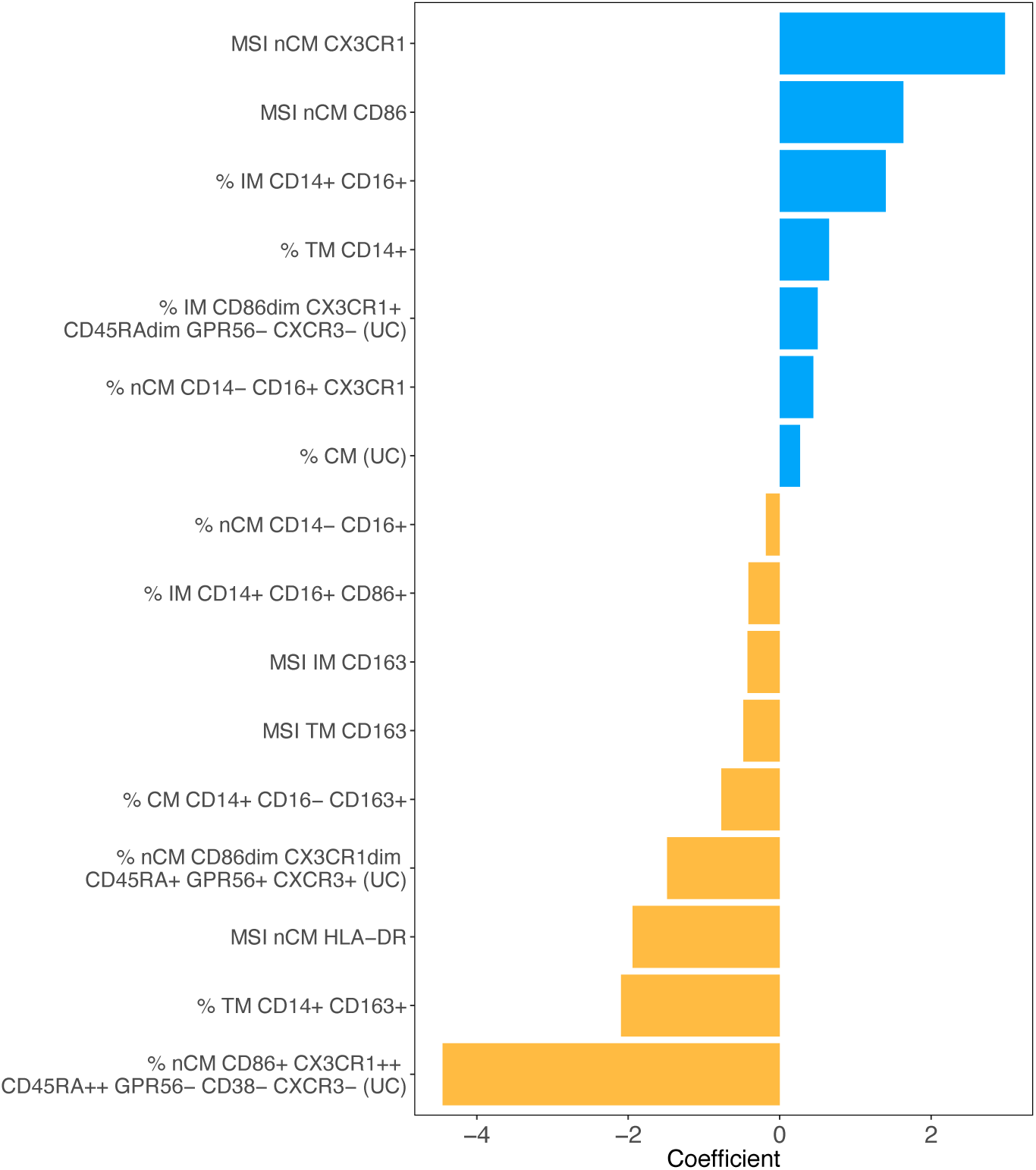
Monocyte features associated with CAD status via regularized regression. Non-zero features obtained from elastic net regression and their coefficients associated with the CAD+/- status. The coefficients were obtained after testing alpha values from 0 to 1 in 0.1 increments. The alpha value that led to the smallest MSE was 0.6 (elastic net regularization). Manual gating and unsupervised clustering variables were used as features. (UC) indicates that the variable was obtained from unsupervised clustering. A total of 16 variables were retained as important predictors of the CAD status after regularization. Seven and nine variables were positively and negatively associated, respectively.

The top 3 variables that were associated with CAD+ were the MSI of CX3CR1 and CD86 in nCM, and the % of IM CD14+ CD16+. The top 3 that were associated with CAD-were the % of nCM CD86+ CX3CR1++ CD45RA++ GPR56- CD38_ CXCR3-identified in the unsupervised clustering, the % of TM CD14+ CD163+, and the MSI of HLA-DR in nCM. The TM CD14+ CD163 agrees with the results obtained from the unsupervised clustering shown in Figure 2.

## DISCUSSION

In this study, mass cytometry and advanced statistical analysis were employed to perform immunophenotyping of monocyte populations and study their associations with infectious and inflammatory diseases, specifically HIV, tuberculosis, and cardiovascular disease. Elastic net regression to identify key features that are most strongly correlated with CAD-/+ status was also applied.

The analysis of the manually gated data showed a statistically significant difference across the CAD/SIS groups regarding the median signal intensity of CD163 in nCM (Figure S5). CD163 has previously been shown to be a potential predictor of mortality and heart failure in people with CAD.^40^ Previous studies have indicated that CD163 is characteristic of anti-inflammatory monocytes and has been identified as marker related to CAD.^41^ The downregulation of CD163 MSI in nCMs among individuals with CAD may occur under specific conditions. For instance, inflammation could promote CD163 shedding producing sCD163 while reducing detectable CD163 surface expression.^42^ Furthermore, in in vitro studies, CD163 downregulation in monocytes and macrophages has been associated to inflammation mediators such as TNF-α, IFN-γ, and LPS.^43^

When comparing the populations obtained from the unsupervised clustering across the CAD/SIS groups, the nCM population CD86dim CX3CR1dim CD45RA+ GPR56+ CXCR3+ was basically absent in the CAD+ >min group. CD86 is a co-stimulatory molecule and plays a crucial role in T cell activation by binding to CD28 on T cells, providing the necessary second signal for full T cell activation. CD86 expression suggests that the subset might be involved in interaction with T cells, likely promoting an immune response.^44^ CX3CR1 is a chemokine receptor predominantly expressed on non-classical monocytes, which actively patrol the vascular endothelium under both homeostatic and inflammatory conditions.^45^ CD45RA is a characteristic marker of naïve T cells and is less well-characterized in monocyte populations.^46^ GPR56 is an adhesion protein coupled-receptor that primarily functions as an inhibitor in natural killer cells. Studies involving GPR56+ monocytes are limited. However, due the similarities between GPR56 and EMR2, it is likely that the function of GPR56+ is similar to monocytes EMR2+ that are involved in the activation and migration of innate cells during systemic inflammation responses.^47^ CXCR3 is another chemokine receptor present in monocytes that can infiltrate CAD-related lesions. CXCR3+ monocytes are associated with plaque instability and adverse cardiovascular events.^48^ It is possible that the lower percentage of this nCM subpopulation among people with CAD >min might indicate that most of these cells may have already migrated to the tissues and are no longer in the bloodstream, thus contributing to atherosclerotic lesion development. Future studies that investigate monocyte and monocyte-derived macrophage subsets both in circulation and atherosclerotic lesions are needed to define the dynamics of this nCM subpopulation and their potential pathogenic role in CAD.

Across the TB/HIV groups, the HIV-TBI+ group exhibited a greater percentage of nCM co-expressing CD86+ and CX3CR1++ compared to the other groups. The CX3CR1 upregulation suggests enhanced chemotactic potential and tissue patrolling capacity, possibly reflecting a more activated, vigilant monocyte phenotype even in latent infection.^6^ Further, a recent study in mice infected with TB showed that the absence of chemokine receptors CX3CR1 was associated with an altered positioning of derived dendritic cells in mediastinal lymph nodes.^49^ The expression of CD86 suggests a role of this population in antigen presentation. Previously it has been reported that circulating non-classical monocytes from HIV-TBI+ individuals exhibit a higher production of IL-6 and TNF-α pro-inflammatory cytokines, compared to HIV-TBI-control individuals.^6^ Future studies can define the functional capacity of this newly classified CD86+ CX3CR1++ CXCR3+ nCM subpopulation and their role in TB infection control and persistent inflammation.

The non-zero features obtained in the elastic net regression provided information about the relevant features associated with the CAD-/+ status. For instance, among the variables strongly associated with CAD+ status, the MSI of CX3XR1 in nCM had the coefficient with the greatest absolute value, which is an indication of a strong immune response and a sign of inflammation in HIV/TB coinfections.^39^ The next positive coefficient was the MSI of CD86 in nCM. It has been shown that expression of CD86 increases as the CAD disease develops.^50^ The CD86+ nCM are potentially geared toward antigen presentation and T-cell stimulation.^51^ The percentage of IM was also positively associated with CAD+. IM are pro-inflammatory and atherogenic.^51^ Also, IM are associated with increased carotid intima-media thickness, a marker of atherosclerosis, and predicted cardiovascular risk even in general population cohorts.^52^ Among the features that were identified as important predictors of CAD-, the nCM population CD86+ CX3CR1++ CD45RA++ GPR56-CD38-CXCR3-showed the greatest negative coefficient. Moderate CD86 expression in non-classical monocytes suggests a role in immune surveillance, rather than driving full activation and inflammation.^53^ In general, CX3CR1 is strongly expressed on non-classical monocytes. CX3CR1+ nCM typically patrol the endothelium and contribute to tissue repair and vascular stability.^45,54^ The other two variables associated with CAD- are the percentage of TM CD14+ CD163+ and the MSI of HLA-DR in nCM. The first of these findings suggests that people with CAD-have higher percentages of CD163+monocytes. These cells are likely anti-inflammatory.^55^ Angiotensin or context-specific studies link high expression of HLA-DR to monocytes with enhanced antigen presentation and immune surveillance traits, especially in non-classical monocytes. One study in HIV populations found that CCR9dim HLA-DR+ nCM were associated with absence of subclinical atherosclerosis, suggesting their HLA-DR+ phenotype is protective.^56^

There are several limitations in this study. For instance, the total number of participants in the sample was small for some of the study groups. Therefore, there might not have been enough power to detect subtle differences in monocyte biomarkers between all groups. Moreover, our cohort includes individuals who live with multiple comorbid conditions, making the analysis even more challenging, a common limitation of human-based clinical and translational studies. The limited sample size does not allow for a detailed analysis or adjustment of all combinations of conditions simultaneously. Still, our identification of specific monocyte subpopulations associated with TB and CAD can be further investigated and validated in future studies. The unsupervised clustering analysis was performed on the manually gated monocyte population, which limited the number of cells that could be included per sample. We randomly selected 10,000 cells per sample or used all cells if the sample contained less than 10,000 cells. This was the case for 15 samples and the minimum number of cells in a sample was 1333. To address potential variability based on the limited sample size, we performed repeated runs of the unsupervised clustering.

The distinct monocyte subpopulations found in this study deserve more study to understand how these populations interact with the diseases as they evolve. This study provides a framework to investigate the functionality of unusual monocyte populations in diseases that affect a large percentage of the human population.

## ACKNOWLEDGEMENTS

Thanks to the study participants.

## Financial support

This work was in part supported by the National Heart, Lung, and Blood Institute (grant number 1R01HL156779 to M.A.H., K23 HL123341 to C.T.L., K01 HL147723 to T.M.T., K23 HL156759 to C.N.W.), the National Center for Advancing Translational Sciences (R03TR004097 to M.A.H), and the National Institute of Arthritis and Musculoskeletal and Skin Diseases (P30AR070549 to S.A.) at the National Institutes of Health, Doris Duke CSDA 2021193 (CNW), Burroughs Wellcome Fund 1021480 (CNW). The content is solely the responsibility of the authors and does not necessarily represent the official views of the National Institutes of Health.

## Disclaimer

The contents are solely the responsibility of the authors and do not necessarily represent the official views of the National Institutes of Health or the institutions with which the authors are affiliated. The funding source had no role in the study design; in the collection, analysis, and interpretation of data; in the writing of the report; or in the decision to submit the report for publication.

---

**Figure S1:**
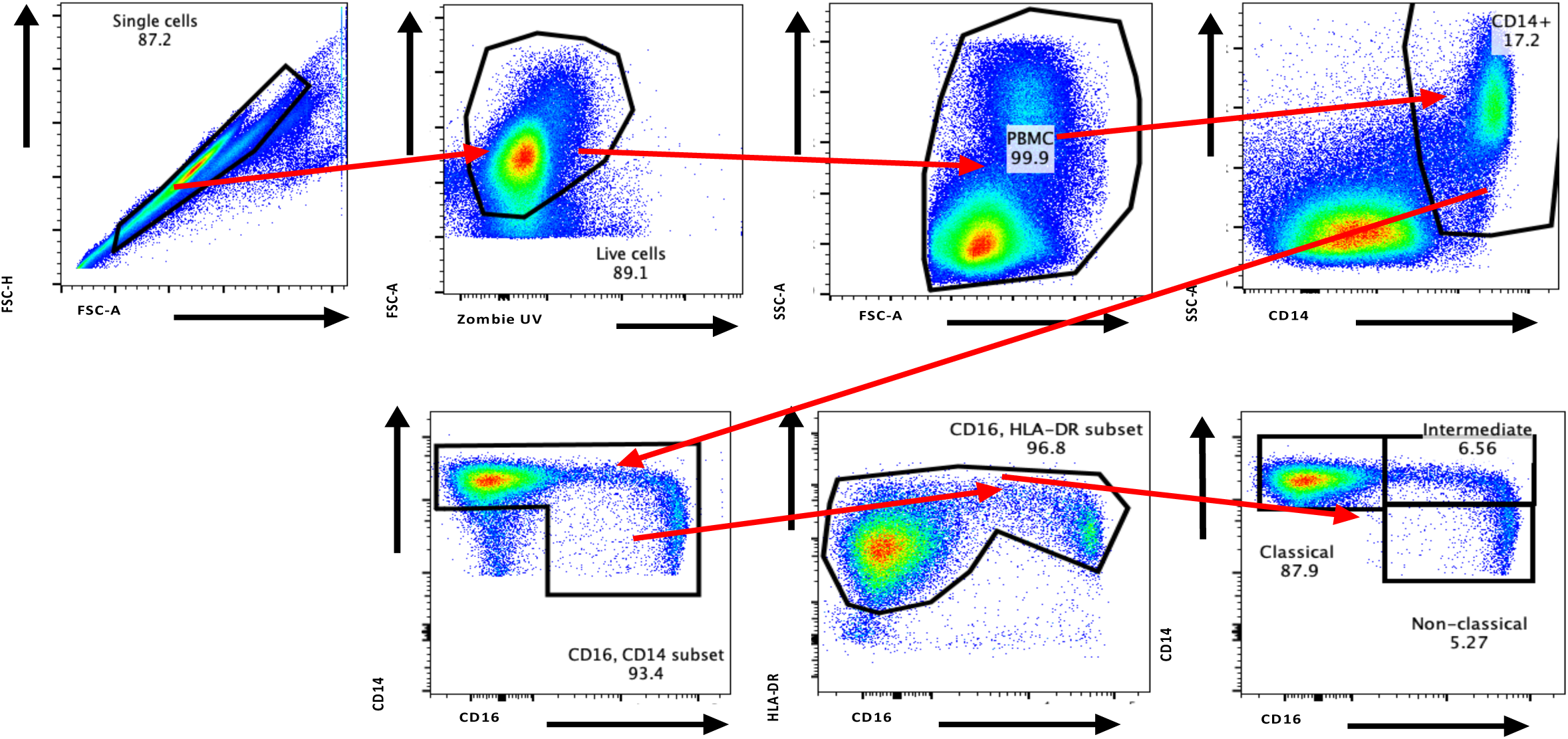
Manual gating strategy sample.

**Figure S2:**
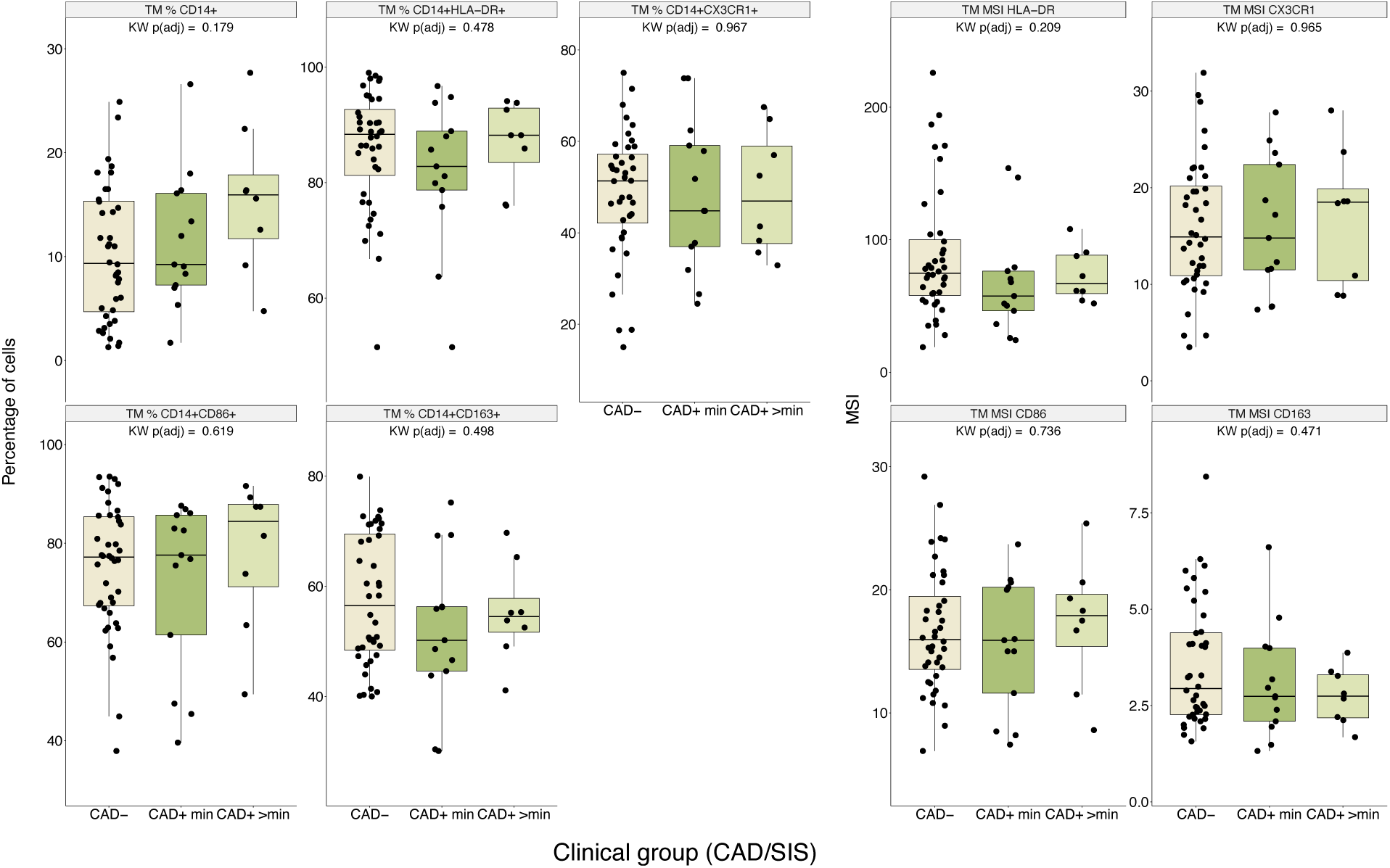
Differences in Total Monocytes Manually Gated Populations and MFI across CAD/SIS. Statistical differences among CAD/SIS study groups across total monocyte (TM) populations and MSI of selected markers. Kruskal-Wallis (KW) tests were performed, and p-values were False Discovery Rate (FDR) adjusted. No statistically significant di_erences were found.

**Figure S3:**
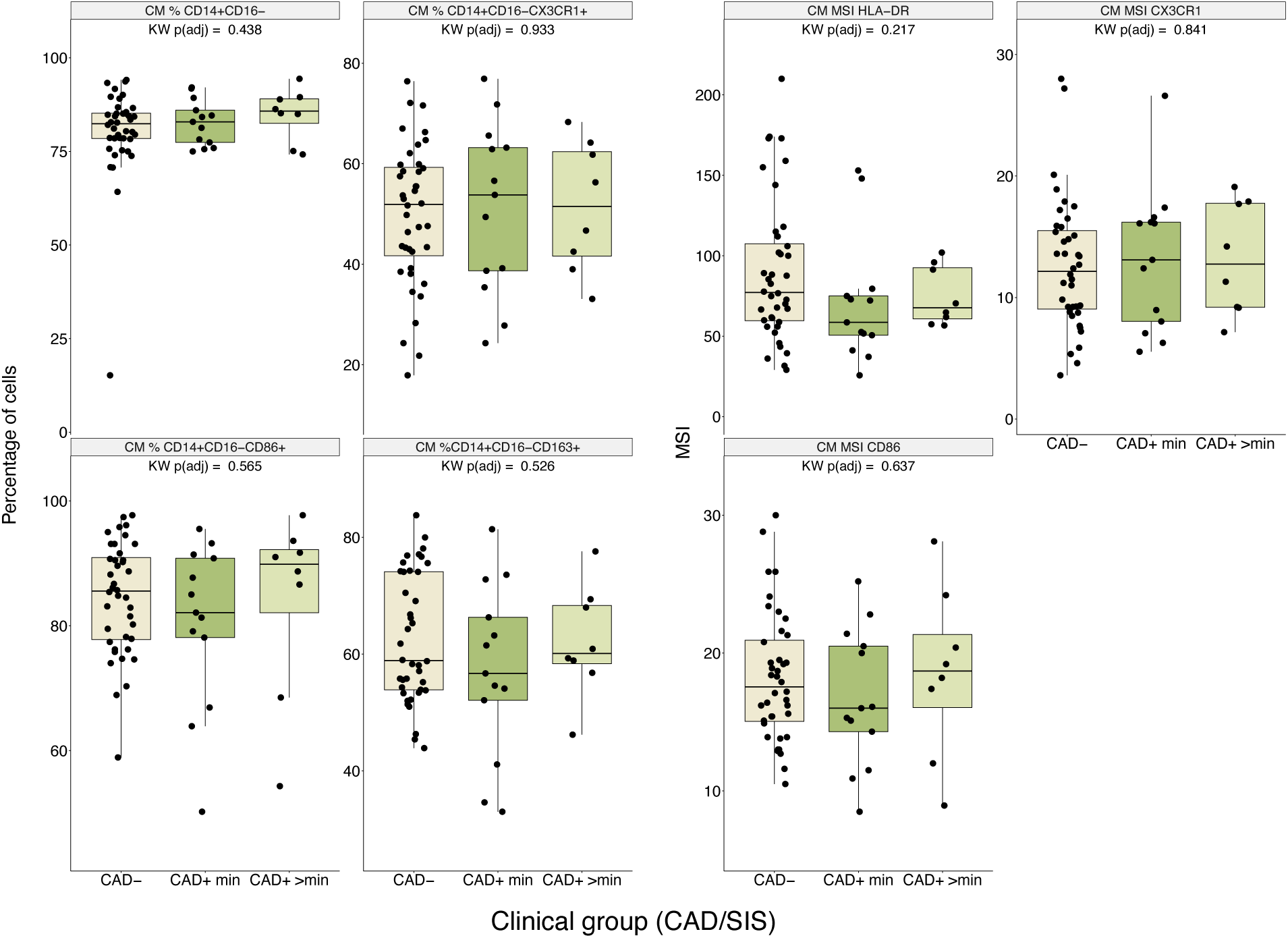
Differences in Classical Monocytes Manually Gated Populations and MFI across CAD/SIS. Statistical differences among CAD/SIS study groups across classical monocyte (CM) populations and MSI of selected markers. Kruskal-Wallis (KW) tests were performed, and p-values were False Discovery Rate (FDR) adjusted. No statistically significant differences were found.

**Figure S4:**
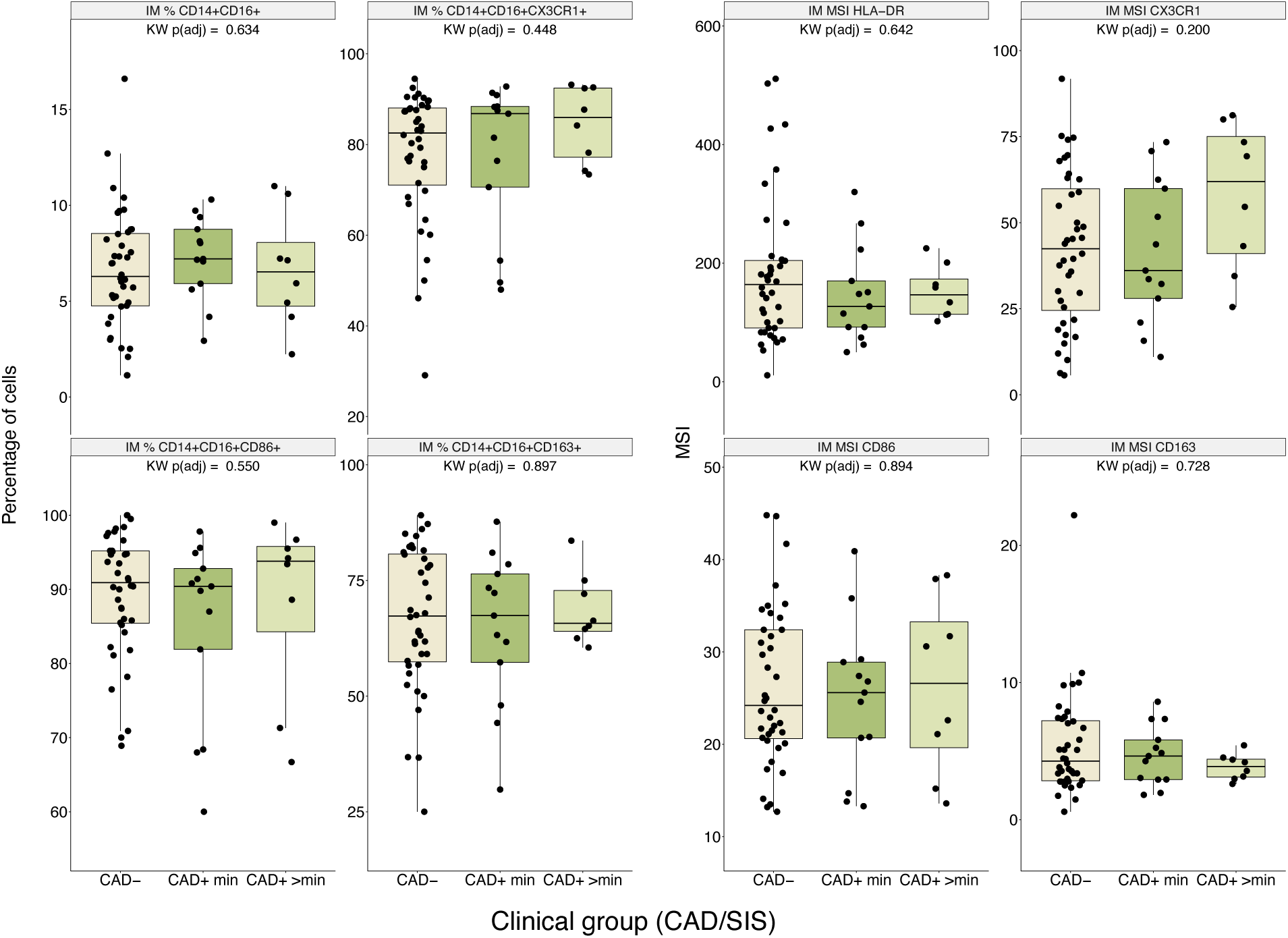
Differences in Intermediate Monocytes Manually Gated Populations and MFI across CAD/SIS. Statistical differences among CAD/SIS study groups across intermediate monocyte (IM) populations and MSI of selected markers. Kruskal-Wallis (KW) tests were performed, and p-values were False Discovery Rate (FDR) adjusted. No statistically significant differences were found.

**Figure S5:**
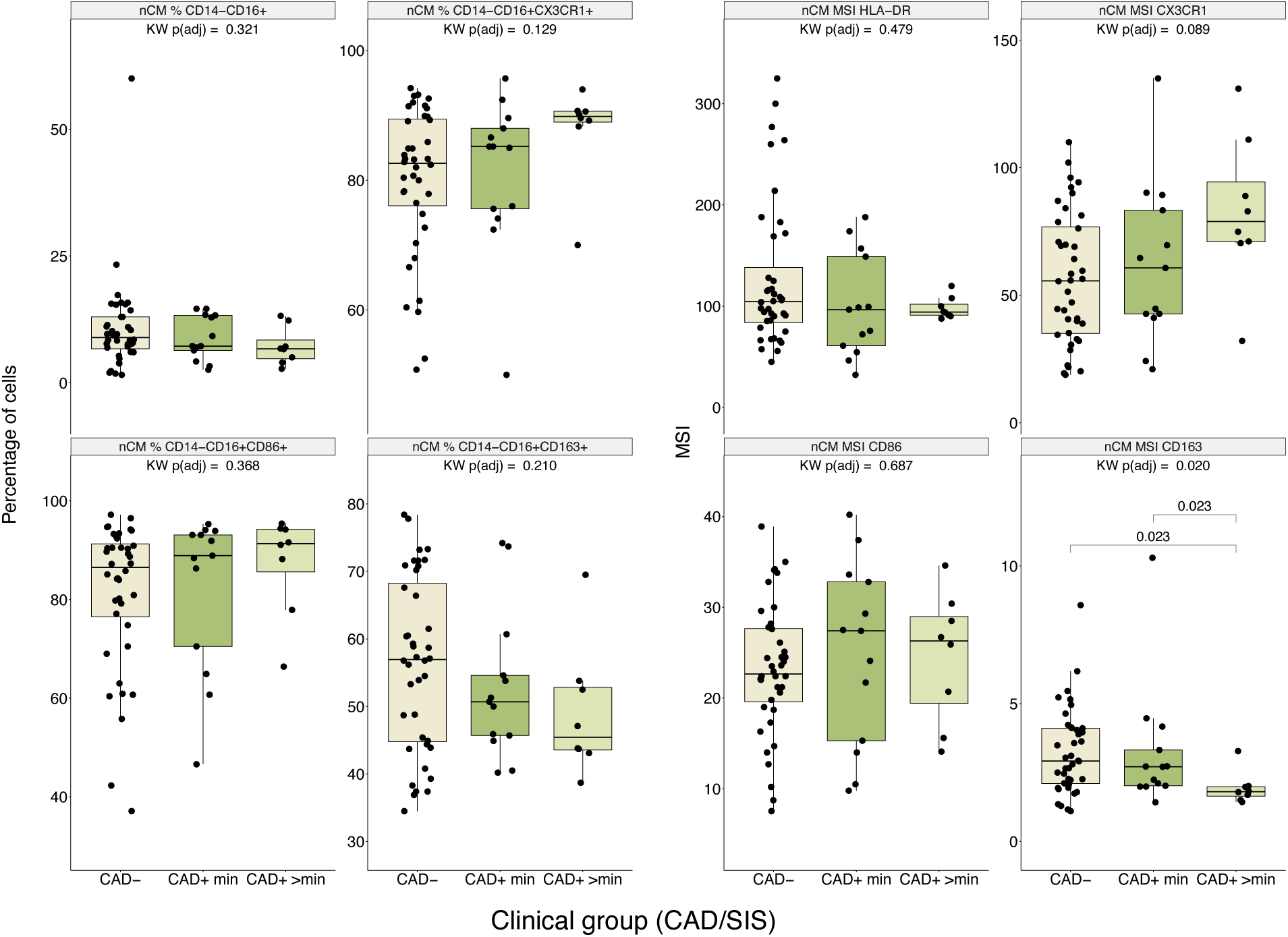
Differences in non Classical Monocytes Manually Gated Populations and MFI across CAD/SIS. Statistical differences among CAD/SIS study groups across non classical monocyte (nCM) populations and MSI of selected markers. Kruskal-Wallis (KW) tests were performed and the p-values adjusted for False Discovery Rate (FDR) using Benjamini-Hochberg method. Only statistically significant (FDR-adjusted p < 0.05) difference was found in the MSI of CD163 in nCM. In post-hoc analyses using Wilcoxon tests, CAD+ >min had significantly lower MSI compared to the other two groups.

**Figure S6:**
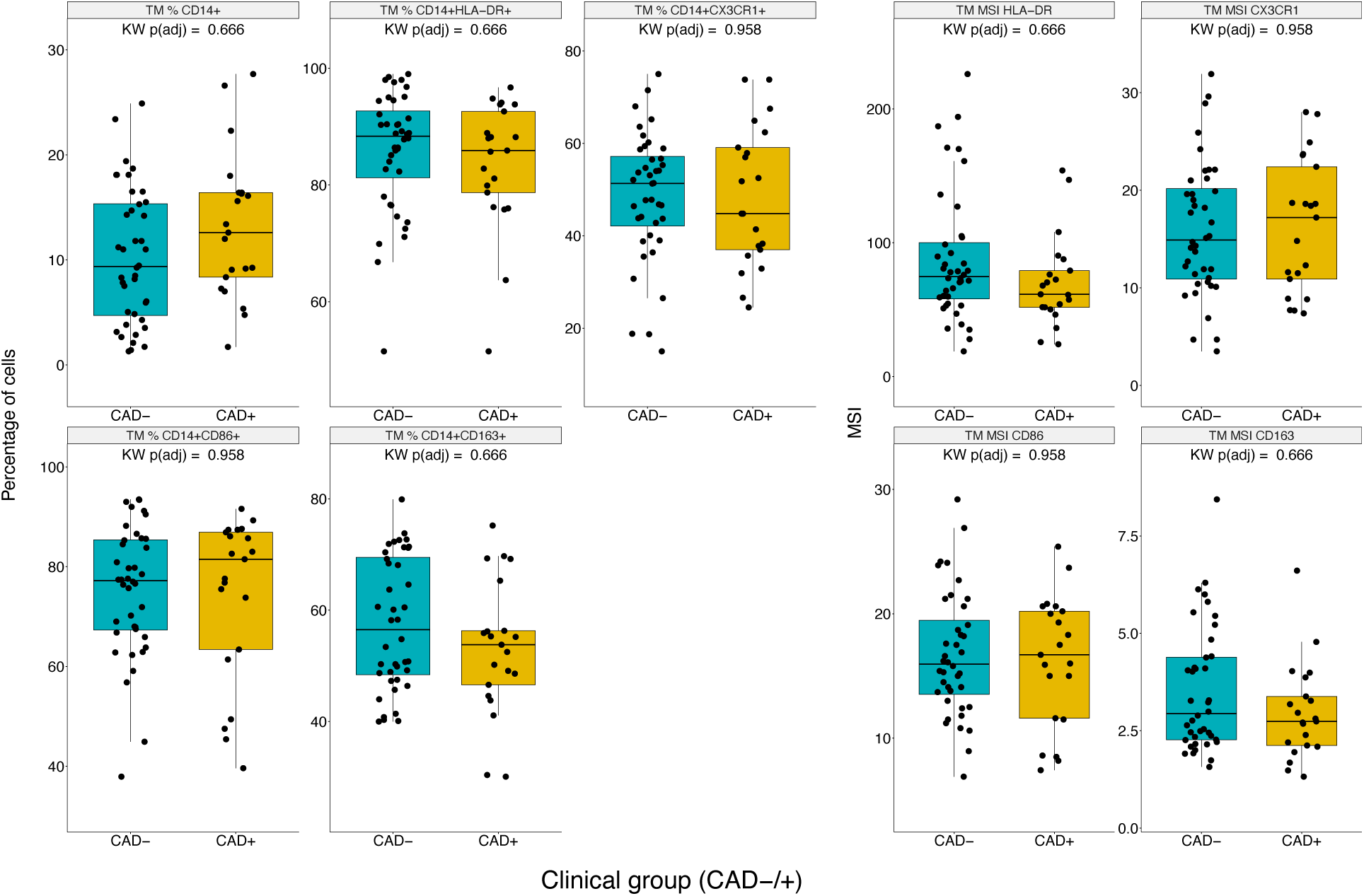
Differences in Total Monocytes Manually Gated Populations and MFI across CAD-/+. Statistical differences between CAD-/+ across total monocytes (TM) populations and MFI of selected markers. Kruskal-Wallis (KW) tests were performed, and p-values were for False Discovery Rate adjusted. No statistically significant differences were found.

**Figure S7:**
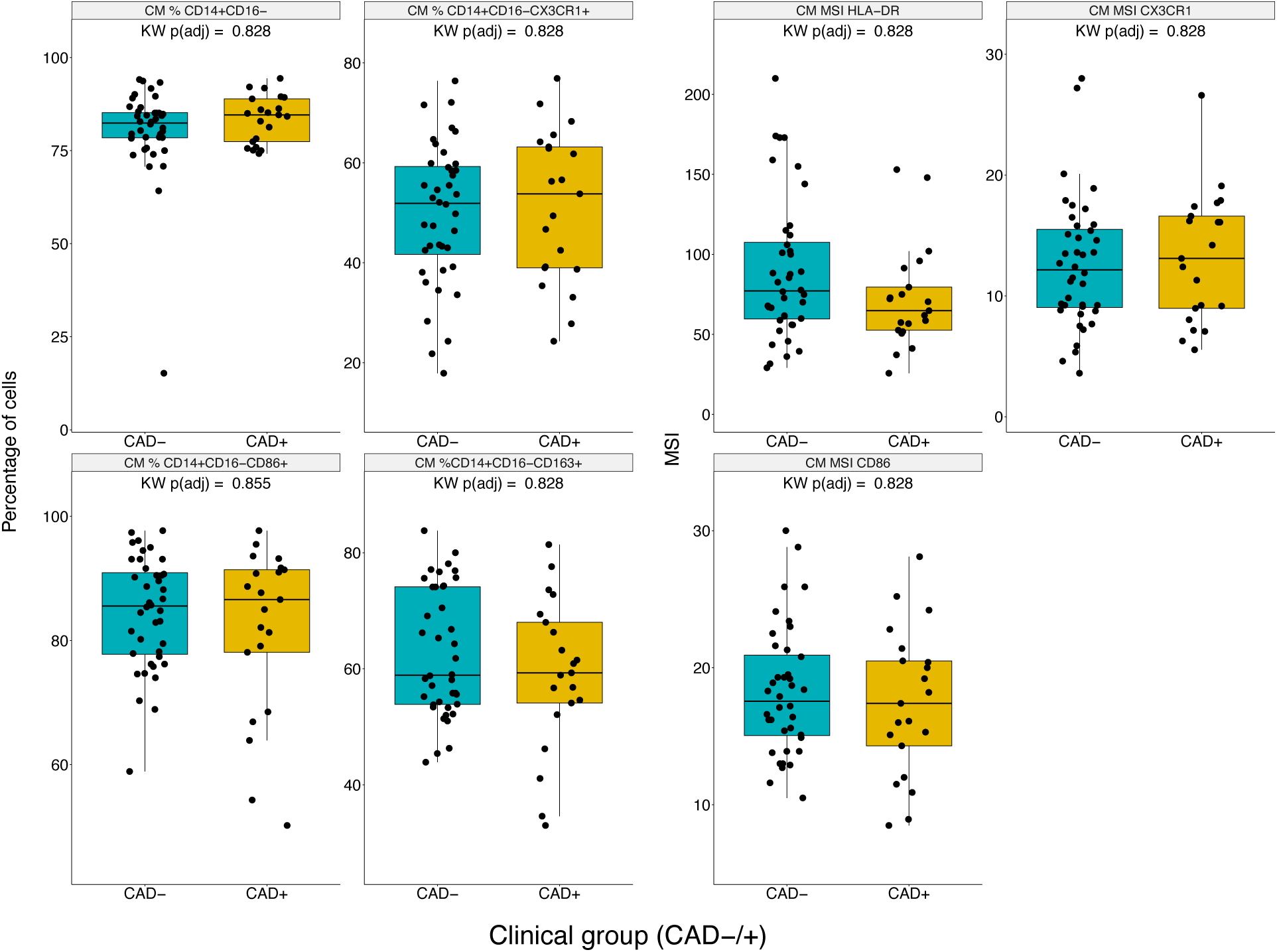
Differences in Classical Monocytes Manually Gated Populations and MFI across CAD-/+. Statistical differences between CAD-/+ across total classical monocyte (CM) populations and MFI of selected markers. Kruskal-Wallis (KW) tests were performed, and p-values were for False Discovery Rate adjusted. No statistically significant differences were found.

**Figure S8:**
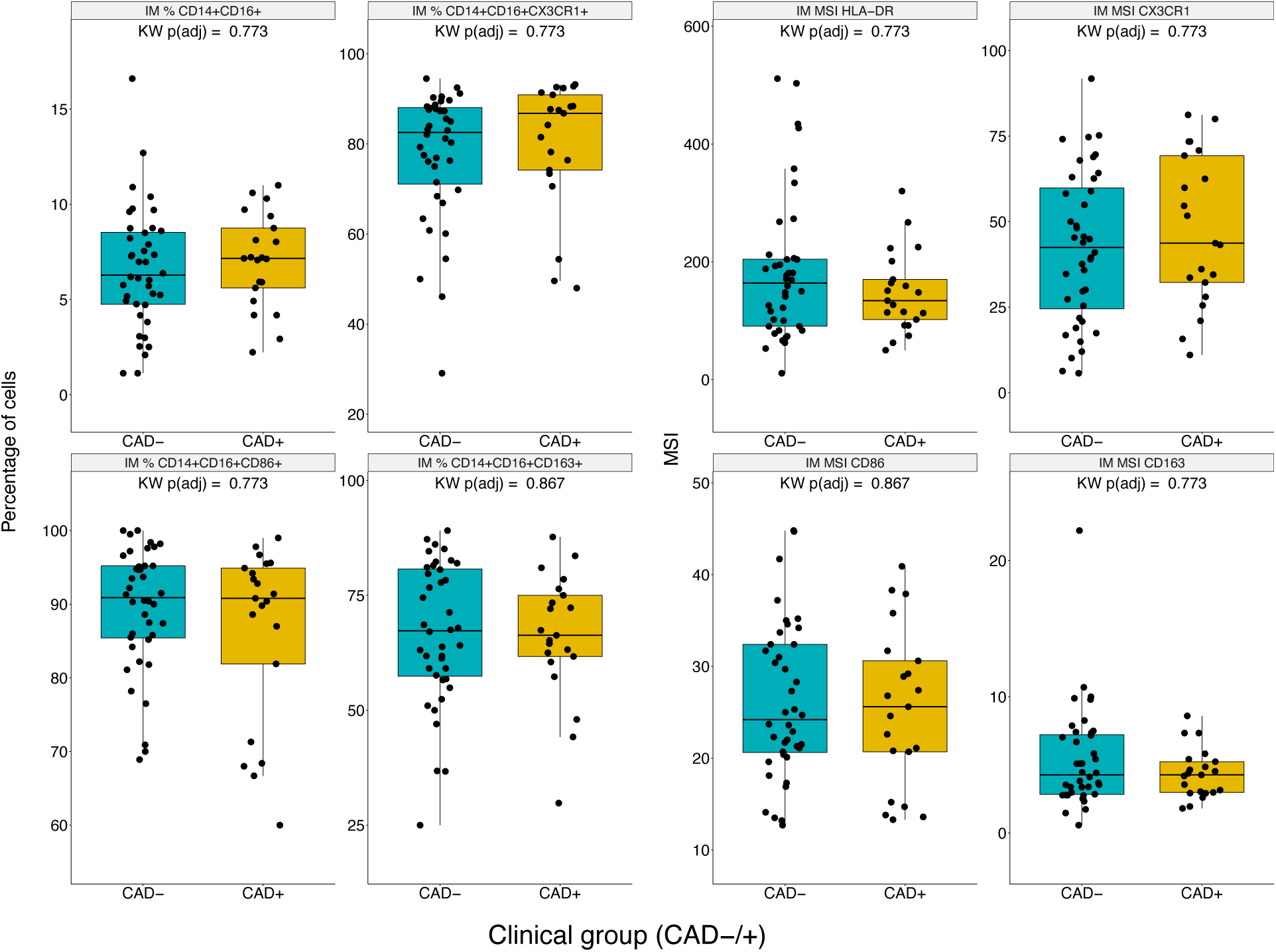
Differences in Intermediate Monocytes Manually Gated Populations and MFI across CAD-/+. Statistical differences between CAD-/+ across intermediate monocyte (IM) populations and MFI of selected markers. Kruskal-Wallis (KW) tests were performed, and p-values were for False Discovery Rate (FDR) adjusted. No statistically significant differences were found.

**Figure S9:**
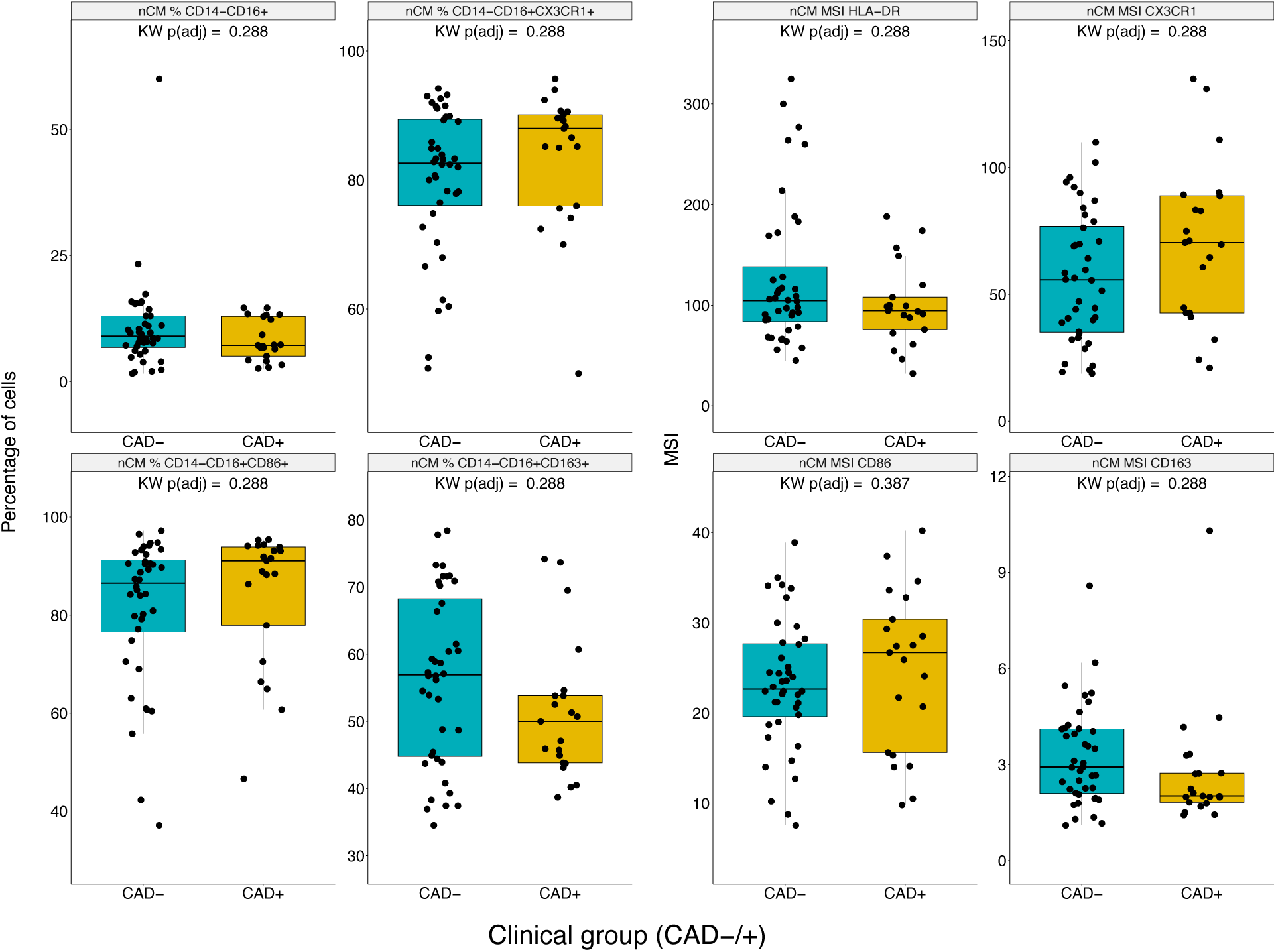
Differences in non Classical Monocytes Manually Gated Populations and MFI across CAD-/+. Statistical differences between CAD-/+ across non classical monocyte (nCM) populations and MFI of selected markers. Kruskal-Wallis (KW) tests were performed, and p-values were False Discovery Rate (FDR) adjusted. No statistically significant differences were found (all FDR-adjusted p ≥ 0.05).

**Figure S10:**
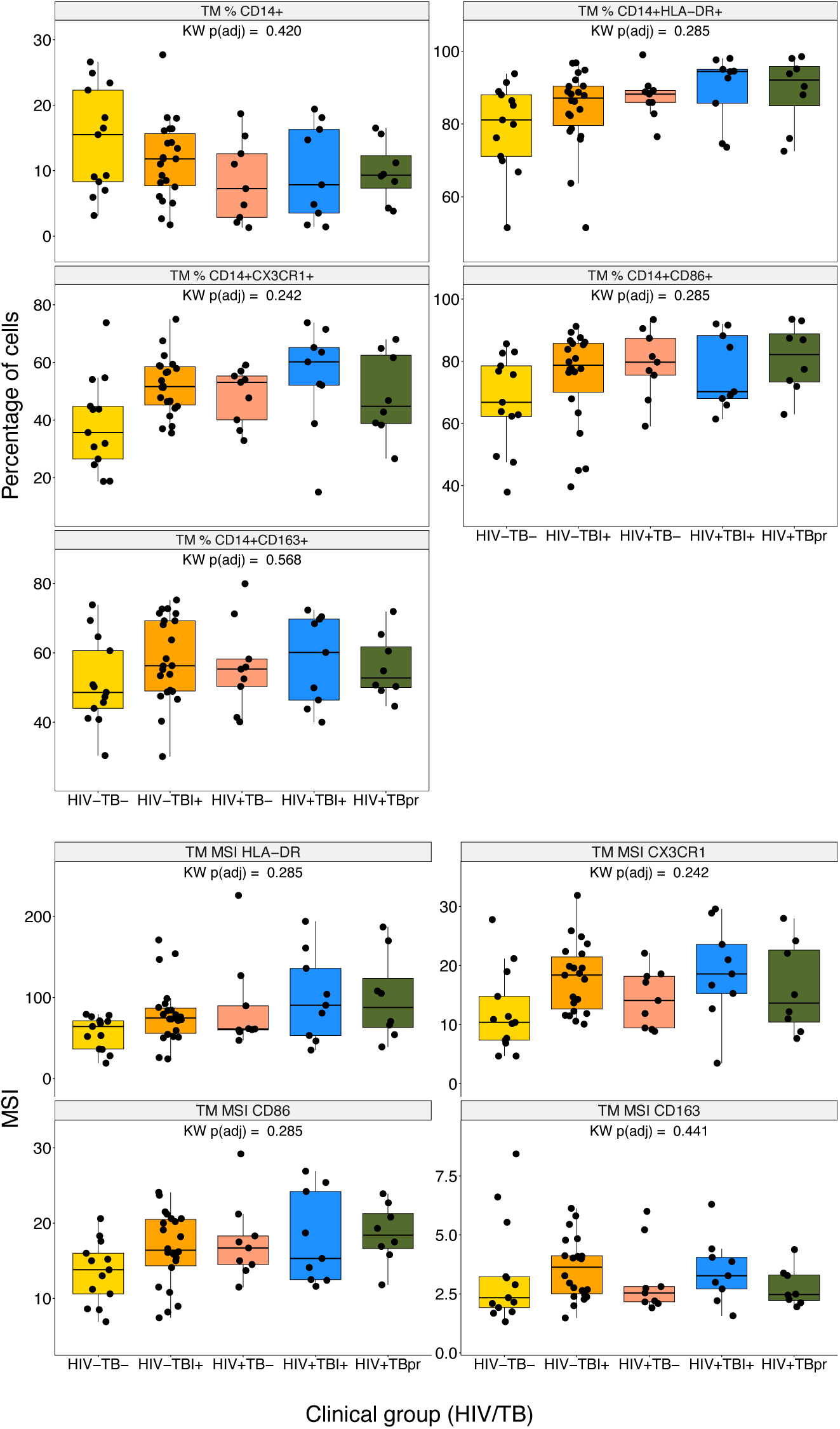
Differences in Total Monocytes Manually Gated Populations and MFI across HIV/TB. Statistical differences among HIV/TB study groups across total monocyte (TM) populations and MSI of selected markers. Kruskal-Wallis (KW) tests were performed, and p-values were False Discovery Rate (FDR) adjusted. No statistically significant differences were found.

**Figure S11:**
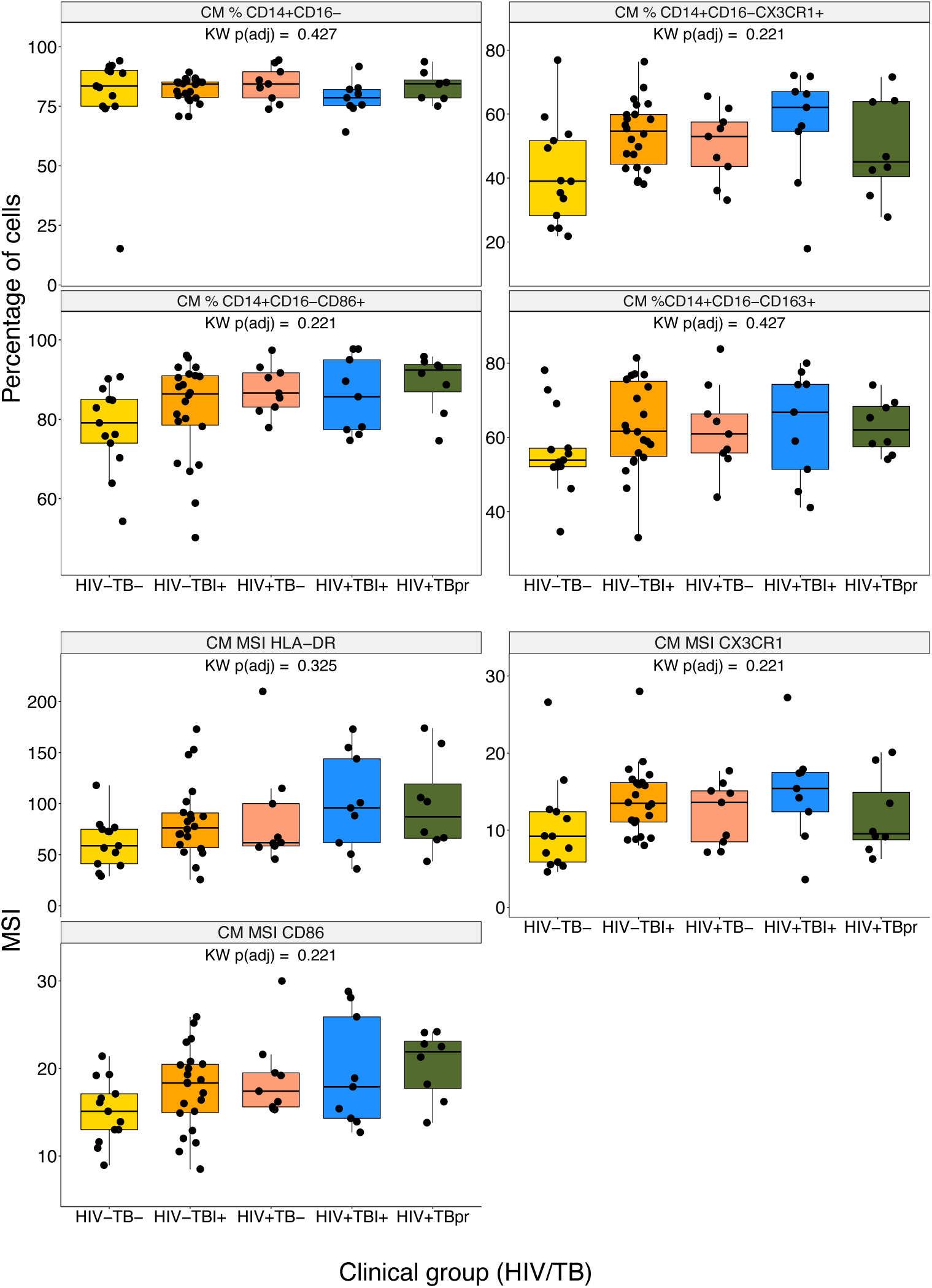
Differences in Classical Monocytes Manually Gated Populations and MFI across HIV/TB. Statistical differences among HIV/TB across classical monocyte (CM) populations and MSI of selected markers. Kruskal-Wallis (KW) tests were performed, and p-values were False Discovery Rate (FDR) adjusted. No statistically significant differences were found.

**Figure S12:**
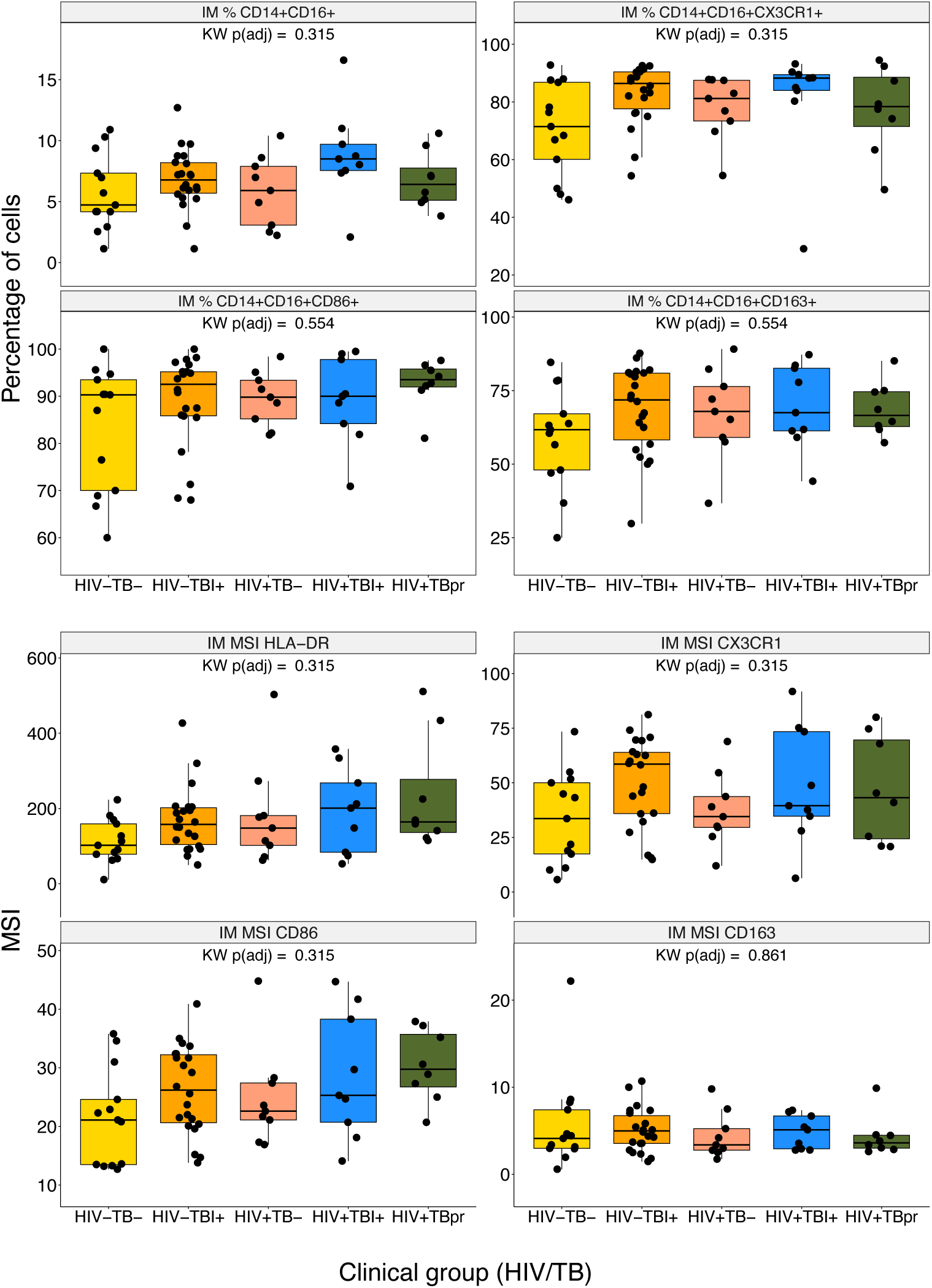
Differences in Intermediate Monocytes Manually Gated Populations and MFI across HIV/TB. Statistical differences among HIV/TB across intermediate monocyte (IM) populations and MSI of selected markers. Kruskal-Wallis (KW) tests were performed, and p-values were False Discovery Rate (FDR) adjusted. No statistically significant differences were found.

**Figure S13:**
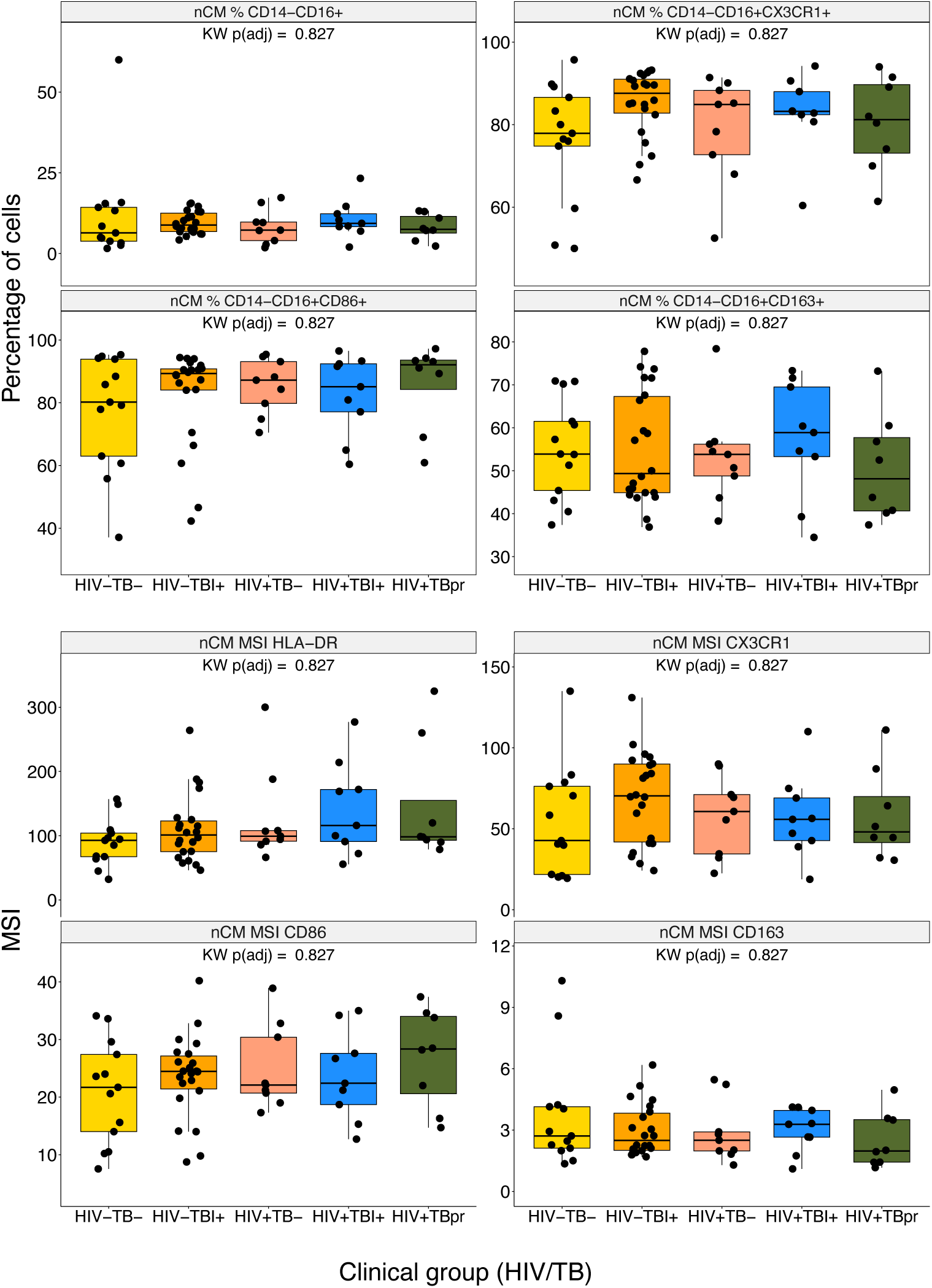
Differences in non Classical Monocytes Manually Gated Populations and MFI across HIV/TB. Statistical differences among HIV/TB across non classical monocyte (nCM) populations and MSI of selected markers. Kruskal-Wallis (KW) tests were performed, and p-values were False Discovery Rate (FDR) adjusted. No statistically significant differences were found.

**Figure S14:**
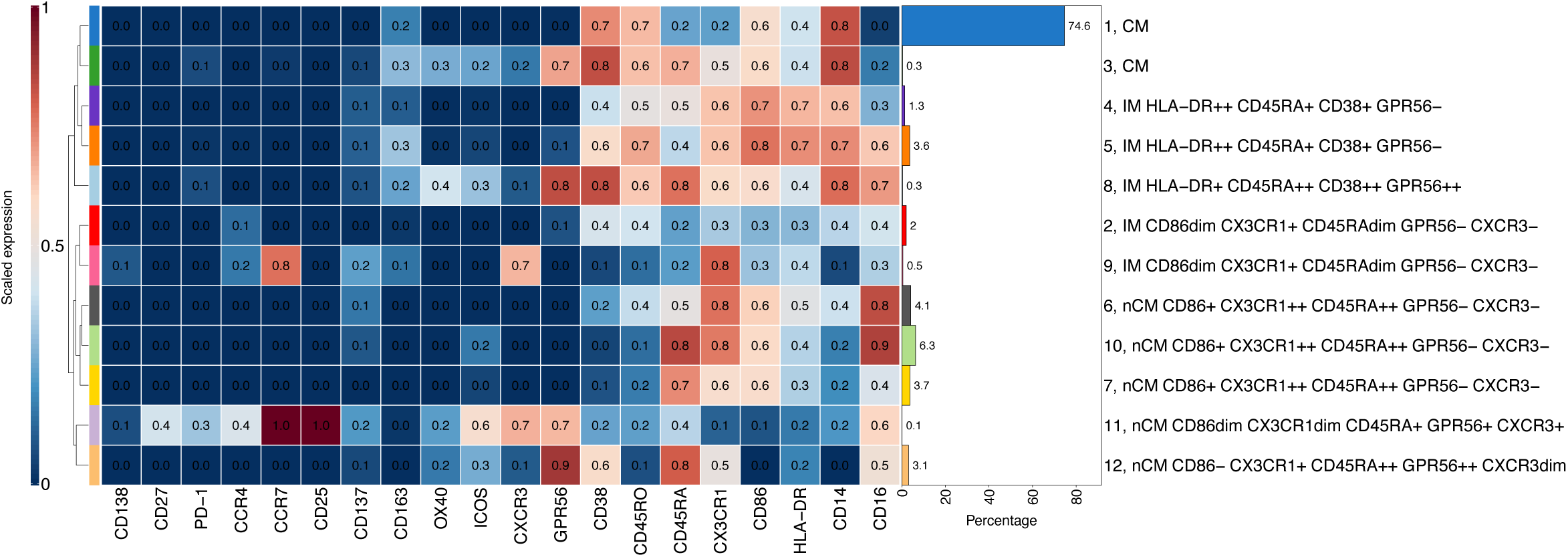
Scaled Median Marker Expressions of the Monocyte Populations Found Using FlowSOM. Based on CD14 and CD16 expression, the 12 populations were classified as 2 classical (CM), 5 intermediate (IM), and 5 non-classical (nCM) monocytes. The bar plot on the right indicates the percentage of the cluster with respect to all cells, i.e. manually gated monocytes, in all the samples. The cluster numbers and marker definitions per cluster are shown on the right. Clusters 1 and 3 (CM), clusters 4 and 5 (IM), clusters 2 and 9 (IM), and clusters 6, 7 and 10 (nCM) were merged due similar characteristics.

**Figure S15:**
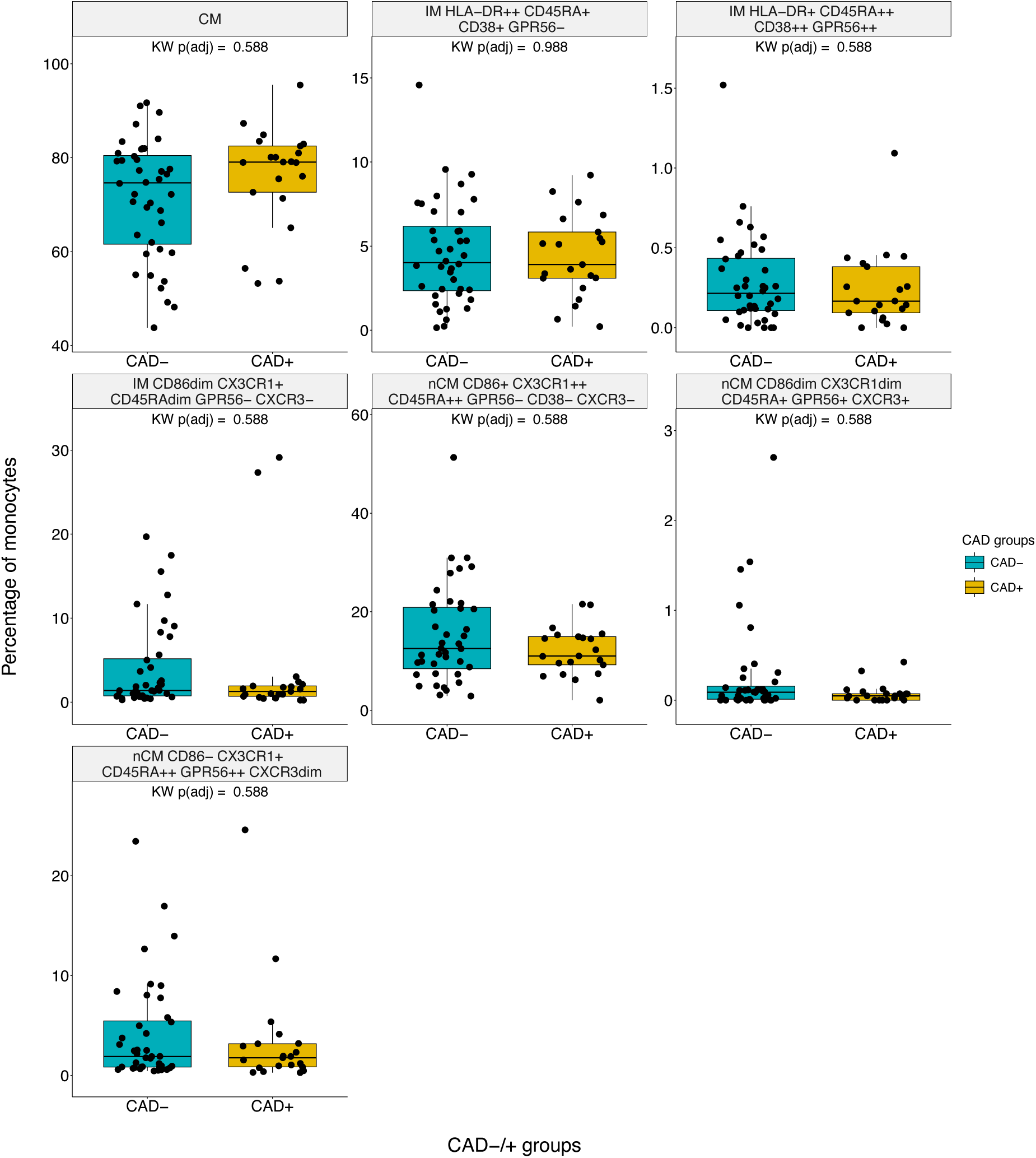
Differences in Monocyte Subset obtained with FlowSOM Across CAD-/+ Groups. Statistical differences of percentages of cells for each monocyte population obtained from unsupervised clustering by FlowSOM across the three CAD-/+ study groups. Kruskal-Wallis (KW) tests were used as omnibus test and p-values were False Discovery Rate (FDR) adjusted. No statistical differences were found all FDR-adjusted p ≥ 0.05.

**Figure S16:**
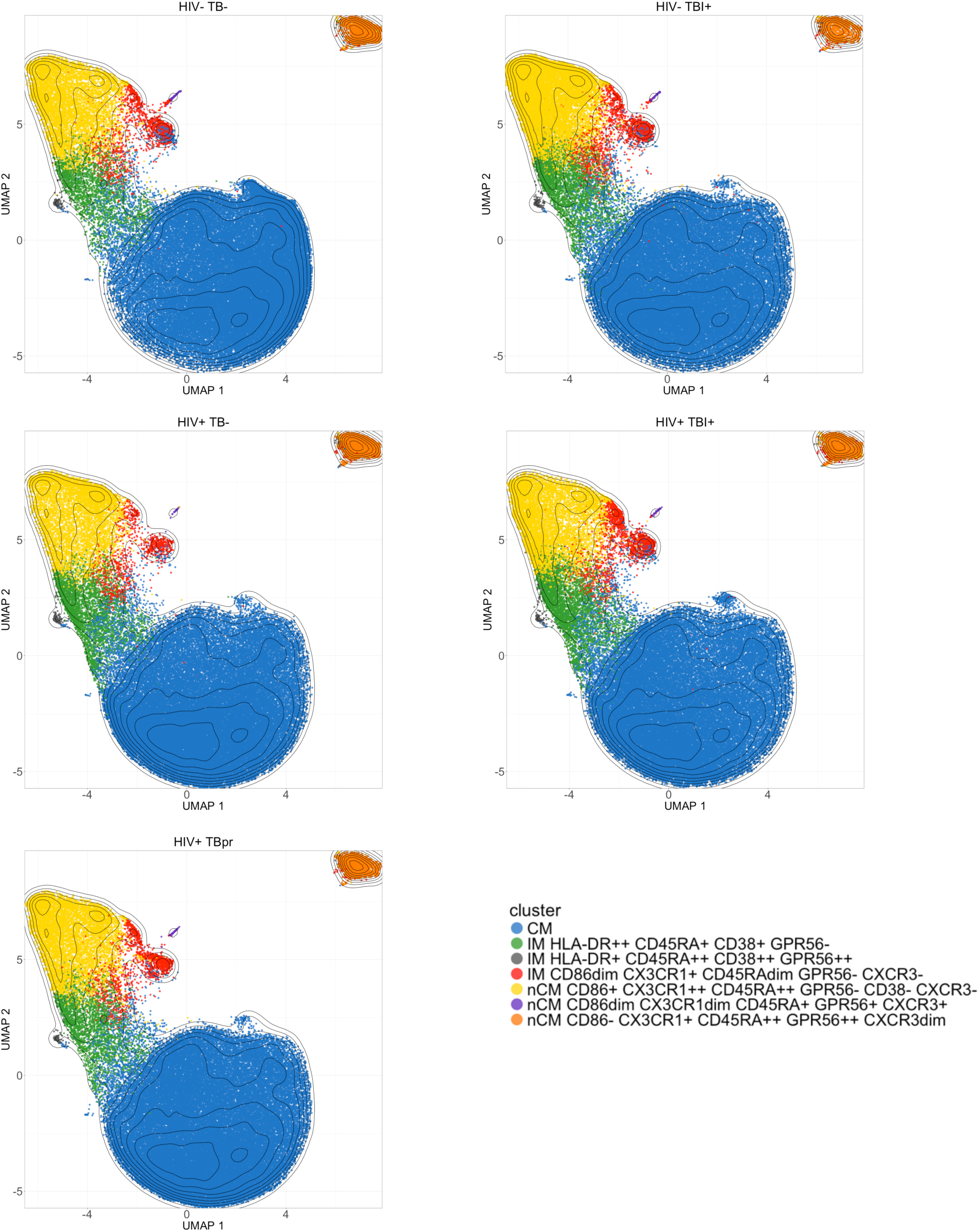
Uniform Manifold Approximation and Projections (UMAPs) stratified by HIV/TB clinical groups. 80000 cells were randomly selected per TB/HIV group and clusters from the FlowSOM analysis were overlaid. Contour lines are of all cells sampled. Colors of the populations match the ones from the bar plot in Figure 1 in the main text.

**Figure S17:**
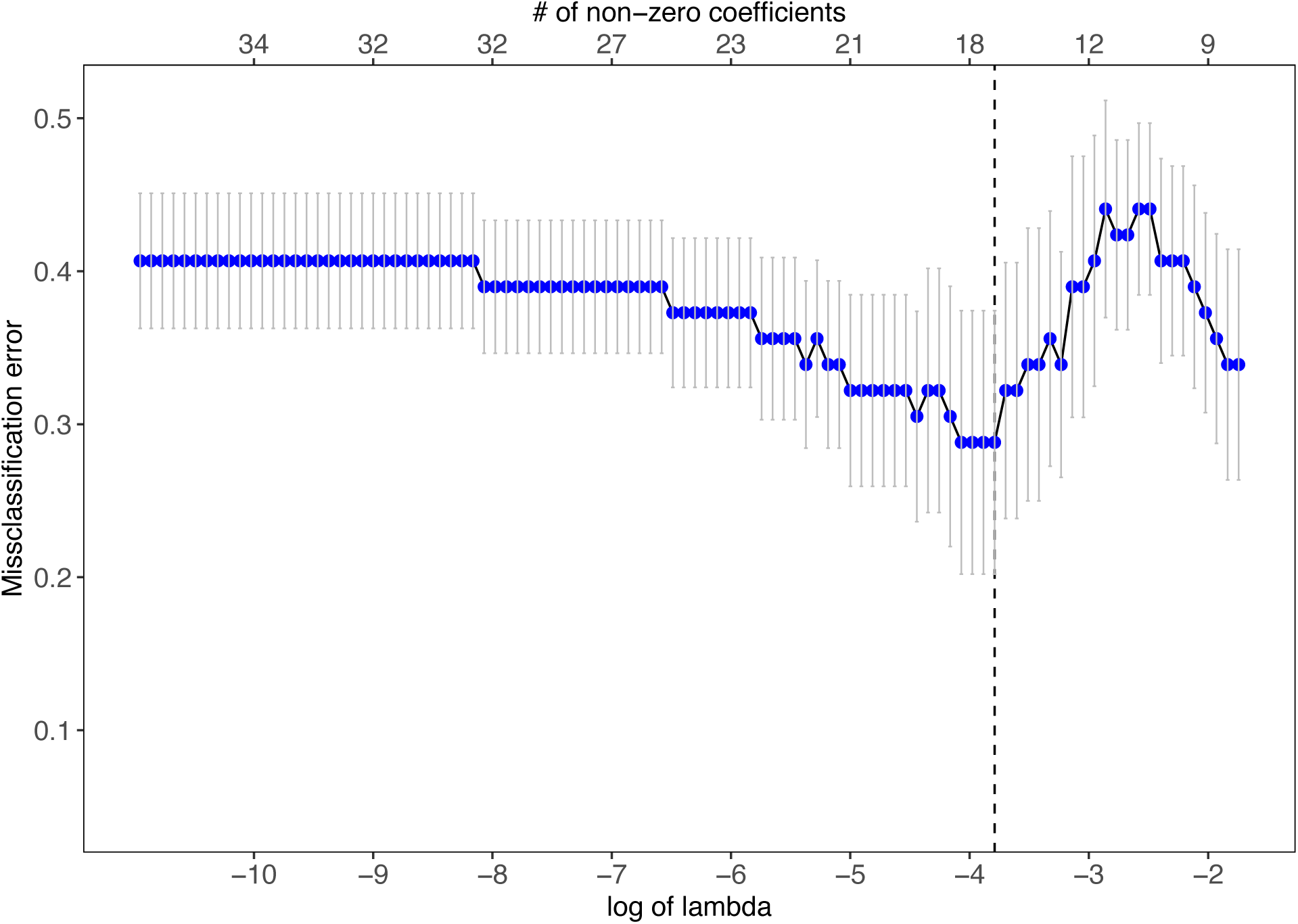
Misclassification error as a function of the log of lambda value (bottom x-axis), and the number of non-zero coefficients (upper x-axis). Dashed line represents the log of lambda that yields the smallest Misclassification error when alpha=0.6 (elastic net regularization). The error bars represent the standard error of the 10-fold cross-validated Misclassification error at each value of lambda. A sequence of one hundred values of lambda was tested, generated using the default settings of the glmnet package. A total of eleven values of alpha (0 – 1 in 0.1 increments) were tested. The non-zero coefficients are plotted in Figure 4 in the main text.

**Table S1.**
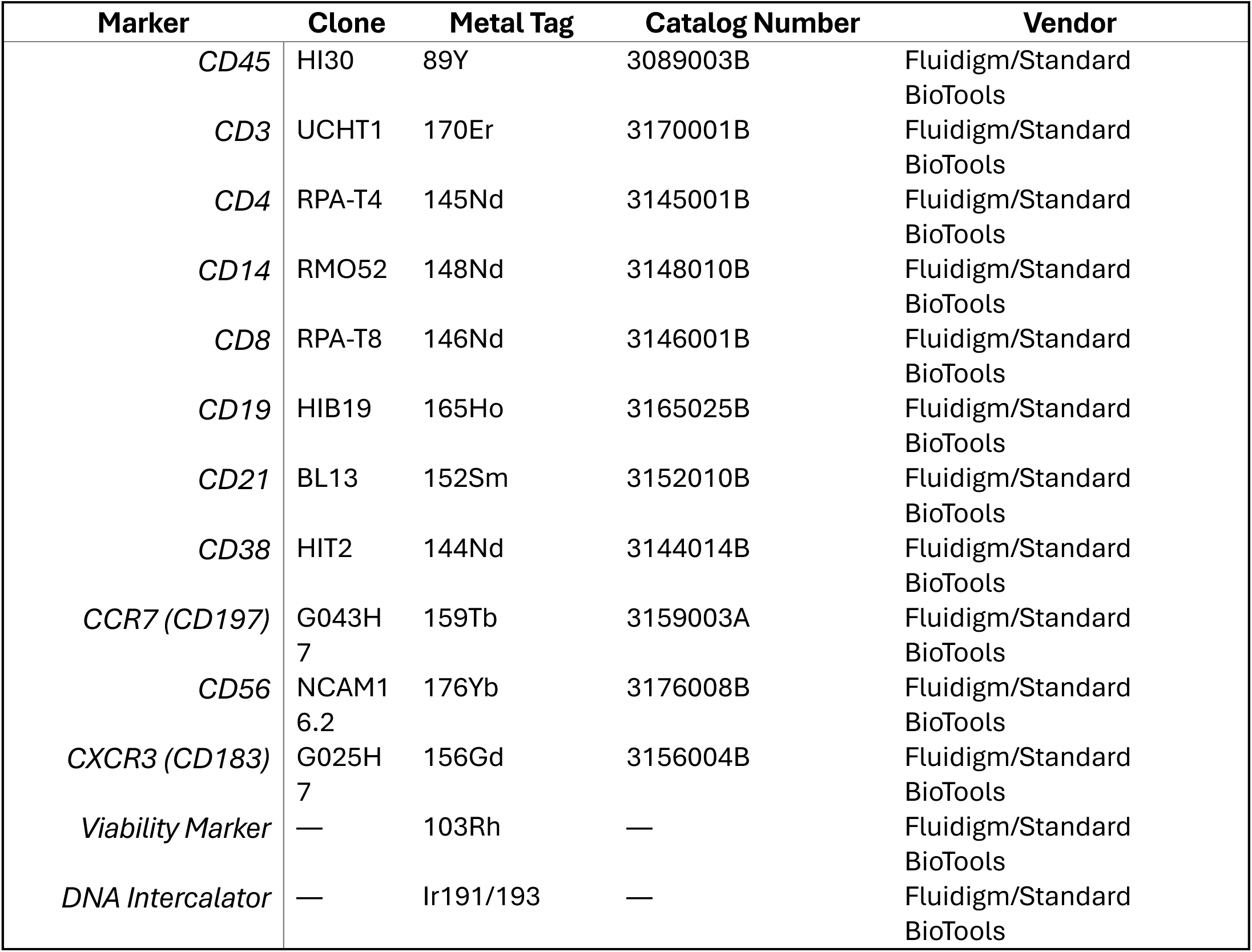
CyTOF Antibody Panel for Immune Cell Phenotyping.

**Table S2:**
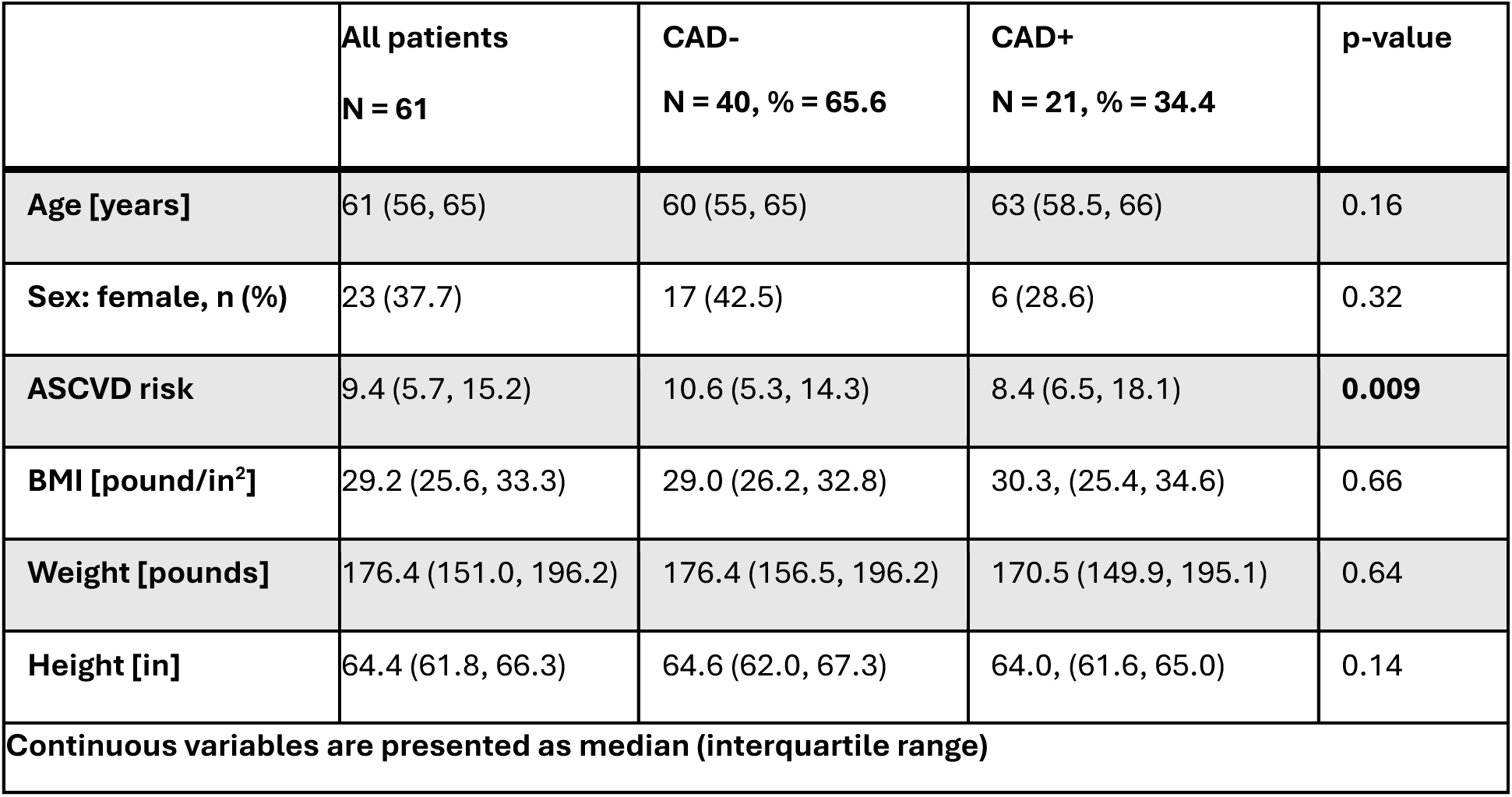
Demographics and clinical characteristics of the cohort using the CAD-/+ classification.

|  | **All patients**<br>**N = 61** | **CAD-**<br>**N = 40, % = 65.6** | **CAD+**<br>**N = 21, % = 34.4** | **p-value** |
| --- | --- | --- | --- | --- |
| **Age [years]** | 61 (56, 65) | 60 (55, 65) | 63 (58.5, 66) | 0.16 |
| **Sex: female, n (%)** | 23 (37.7) | 17 (42.5) | 6 (28.6) | 0.32 |
| **ASCVD risk** | 9.4 (5.7, 15.2) | 10.6 (5.3, 14.3) | 8.4 (6.5, 18.1) | **0.009** |
| **BMI [pound/in<sup>2</sup>]** | 29.2 (25.6, 33.3) | 29.0 (26.2, 32.8) | 30.3, (25.4, 34.6) | 0.66 |
| **Weight [pounds]** | 176.4 (151.0, 196.2) | 176.4 (156.5, 196.2) | 170.5 (149.9, 195.1) | 0.64 |
| **Height [in]** | 64.4 (61.8, 66.3) | 64.6 (62.0, 67.3) | 64.0, (61.6, 65.0) | 0.14 |
| **Continuous variables are presented as median (interquartile range)** |  |  |  |  |

## Notes

### Competing Interest Statement

The authors have declared no competing interest.

## REFERENCES

1. Govender RD, Hashim MJ, Khan MA, Mustafa H, Khan G. Global Epidemiology of HIV/AIDS: A Resurgence in North America and Europe: JEGH. 2021;11(3):296.

2. Berida T, Lindsley CW. Move over COVID, Tuberculosis Is Once again the Leading Cause of Death from a Single Infectious Disease. J Med Chem. 2024 Dec 26;67(24):21633– 21640.

3. Gaidai O, Cao Y, Loginov S. Global Cardiovascular Diseases Death Rate Prediction. Current Problems in Cardiology. 2023 May 1;48(5):101622.

4. Siedner MJ, Ghoshhajra B, Erem G, Nassanga R, Randhawa M, Ochieng A, Acan M, Lu MT, Thondapu V, Takigami A, Reynolds Z, Atwiine F, Tindimwebwa E, Gilbert RF, Passell E, Sagar S, Tong Y, Ntusi NAB, Tsai AC, Bibangambah P, Gaziano T, Hoeppner SS, Longenecker CT, Okello S, Asiimwe S. Epidemiology of Coronary Atherosclerosis Among People Living With HIV in Uganda. Ann Intern Med. American College of Physicians; 2025 Apr 15;178(4):468–478.

5. Hyle EP, Mayosi BM, Middelkoop K, Mosepele M, Martey EB, Walensky RP, Bekker LG, Triant VA. The association between HIV and atherosclerotic cardiovascular disease in sub-Saharan Africa: a systematic review. BMC Public Health. 2017 Dec;17(1):954.

6. Feria MG, Chang C, Ticona E, Moussa A, Zhang B, Ballena I, Azañero R, Ticona C, De Cecco CN, Fichtenbaum CJ, O’Donnell RE, La Rosa A, Sanchez J, Andorf S, Atehortua L, Katz JD, Chougnet CA, Deepe GS Jr, Huaman MA. Pro-Inflammatory Alterations of Circulating Monocytes in Latent Tuberculosis Infection. Open Forum Infectious Diseases. 2022 Dec 1;9(12):ofac629.

7. Okello S, Amir A, Bloomfield GS, Kentoffio K, Lugobe HM, Reynolds Z, Magodoro IM, North CM, Okello E, Peck R, Siedner MJ. Prevention of cardiovascular disease among people living with HIV in sub-Saharan Africa. Progress in Cardiovascular Diseases. 2020 Mar;63(2):149–159.

8. Obare LM, Temu T, Mallal SA, Wanjalla CN. Inflammation in HIV and Its Impact on Atherosclerotic Cardiovascular Disease. Circulation Research. 2024 May 24;134(11):1515– 1545.

9. Rambaran S, Maseko TG, Lewis L, Hassan-Moosa R, Archary D, Ngcapu S, Garrett N, McKinnon LR, Padayatchi N, Naidoo K, Sivro A. Blood monocyte and dendritic cell profiles among people living with HIV with Mycobacterium tuberculosis co-infection. BMC Immunol. 2023 Jul 21;24(1):21.

10. Huaman MA, Henson D, Ticona E, Sterling TR, Garvy BA. Tuberculosis and cardiovascular disease: linking the epidemics. Trop Dis Travel Med Vaccines. 2015 Dec;1(1):10.

11. Chidambaram V, Ruelas Castillo J, Kumar A, Wei J, Wang S, Majella MG, Gupte A, Wang JY, Karakousis PC. The association of atherosclerotic cardiovascular disease and statin use with inflammation and treatment outcomes in tuberculosis. Sci Rep. 2021 Jul 27;11(1):15283.

12. Knudsen AD, Bouazzi R, Afzal S, Gelpi M, Benfield T, Høgh J, Thomsen MT, Trøseid M, Nordestgaard BG, Nielsen SD. Monocyte count and soluble markers of monocyte activation in people living with HIV and uninfected controls. BMC Infect Dis. 2022 Dec;22(1):451.

13. Wallis ZK, Williams KC. Monocytes in HIV and SIV Infection and Aging: Implications for Inflamm-Aging and Accelerated Aging. Viruses. 2022 Feb 17;14(2):409. PMCID: PMC8880456

14. Lyu M, Xu G, Zhou J, Reboud J, Wang Y, Lai H, Chen Y, Zhou Y, Zhu G, Cooper JM, Ying B. Single-Cell Sequencing Reveals Functional Alterations in Tuberculosis. Advanced Science. 2024;11(11):2305592.

15. Ji H, Li Y, Fan Z, Zuo B, Jian X, Li L, Liu T. Monocyte/lymphocyte ratio predicts the severity of coronary artery disease: a syntax score assessment. BMC Cardiovascular Disorders. 2017 Mar 31;17(1):90.

16. Moroni F, Ammirati E, Norata GD, Magnoni M, Camici PG. The Role of Monocytes and Macrophages in Human Atherosclerosis, Plaque Neoangiogenesis, and Atherothrombosis. Mediators of Inflammation. 2019;2019(1):7434376.

17. Tapp LD, Shantsila E, Wrigley BJ, Pamukcu B, Lip GYH. The CD14++CD16+ monocyte subset and monocyte-platelet interactions in patients with ST-elevation myocardial infarction. Journal of Thrombosis and Haemostasis. Elsevier; 2012 Jul 1;10(7):1231–1241. PMID: 22212813

18. Eligini S, Cosentino N, Fiorelli S, Fabbiocchi F, Niccoli G, Refaat H, Camera M, Calligaris G, De Martini S, Bonomi A, Veglia F, Fracassi F, Crea F, Marenzi G, Tremoli E. Biological profile of monocyte-derived macrophages in coronary heart disease patients: implications for plaque morphology. Sci Rep. Nature Publishing Group; 2019 Jun 18;9(1):8680.

19. Amengual J, Barrett TJ. Monocytes and macrophages in atherogenesis. Curr Opin Lipidol. 2019 Oct;30(5):401–408. PMCID: PMC7809604

20. Sampath P, Moideen K, Ranganathan UD, Bethunaickan R. Monocyte Subsets: Phenotypes and Function in Tuberculosis Infection. Front Immunol [Internet]. Frontiers; 2018 Jul 30 [cited 2025 Feb 18];9. Available from: https://www.frontiersin.org/journals/immunology/articles/10.3389/fimmu.2018.01726/full

21. Ma R, Yang W, Guo W, Zhang H, Wang Z, Ge Z. Single-cell transcriptome analysis reveals the dysregulated monocyte state associated with tuberculosis progression. BMC Infectious Diseases. 2025 Feb 12;25(1):210.

22. Bai R, Li Z, Lv S, Wang R, Hua W, Wu H, Dai L. Persistent Inflammation and Non-AIDS Comorbidities During ART: Coming of the Age of Monocytes. Front Immunol [Internet]. Frontiers; 2022 Apr 11 [cited 2025 Feb 18];13. Available from: https://www.frontiersin.org/journals/immunology/articles/10.3389/fimmu.2022.820480/full

23. Wong ME, Jaworowski A, Hearps AC. The HIV Reservoir in Monocytes and Macrophages. Front Immunol [Internet]. Frontiers; 2019 Jun 26 [cited 2025 Feb 18];10. Available from: https://www.frontiersin.org/journals/immunology/articles/10.3389/fimmu.2019.01435/full

24. Williams DW, Veenstra M, Gaskill PJ, Morgello S, Calderon TM, Berman JW. Monocytes Mediate HIV Neuropathogenesis: Mechanisms that Contribute to HIV Associated Neurocognitive Disorders. Curr HIV Res. 2014;12(2):85–96. PMCID: PMC4351961

25. Ellery PJ, Tippett E, Chiu YL, Paukovics G, Cameron PU, Solomon A, Lewin SR, Gorry PR, Jaworowski A, Greene WC, Sonza S, Crowe SM. The CD16+ Monocyte Subset Is More Permissive to Infection and Preferentially Harbors HIV-1 In Vivo1. The Journal of Immunology. 2007 May 15;178(10):6581–6589.

26. Longenecker CT, Bogorodskaya M, Margevicius S, Nazzinda R, Bittencourt MS, Erem G, Nalukwago S, Huaman MA, Ghoshhajra BB, Siedner MJ, Juchnowski SM, Zidar DA, McComsey GA, Kityo C. Sex modifies the association between HIV and coronary artery disease among older adults in Uganda. Journal of the International AIDS Society. 2022;25(1):e25868.

27. Goff DC, Lloyd-Jones DM, Bennett G, Coady S, D’Agostino RB, Gibbons R, Greenland P, Lackland DT, Levy D, O’Donnell CJ, Robinson JG, Schwartz JS, Shero ST, Smith SC, Sorlie P, Stone NJ, Wilson PWF. 2013 ACC/AHA Guideline on the Assessment of Cardiovascular Risk. Circulation. American Heart Association; 2014 Jun 24;129(25_suppl_2):S49–S73.

28. Min JK, Shaw LJ, Devereux RB, Okin PM, Weinsaft JW, Russo DJ, Lippolis NJ, Berman DS, Callister TQ. Prognostic Value of Multidetector Coronary Computed Tomographic Angiography for Prediction of All-Cause Mortality. JACC. American College of Cardiology Foundation; 2007 Sep 18;50(12):1161–1170.

29. Hahne F, LeMeur N, Brinkman RR, Ellis B, Haaland P, Sarkar D, Spidlen J, Strain E, Gentleman R. flowCore: a Bioconductor package for high throughput flow cytometry. BMC Bioinformatics. 2009 Apr 9;10:106. PMCID: PMC2684747

30. Gentleman RC, Carey VJ, Bates DM, Bolstad B, Dettling M, Dudoit S, Ellis B, Gautier L, Ge Y, Gentry J, Hornik K, Hothorn T, Huber W, Iacus S, Irizarry R, Leisch F, Li C, Maechler M, Rossini AJ, Sawitzki G, Smith C, Smyth G, Tierney L, Yang JY, Zhang J. Bioconductor: open software development for computational biology and bioinformatics. Genome Biol. 2004 Sep 15;5(10):R80.

31. Bendall SC, Simonds EF, Qiu P, Amir E ad D, Krutzik PO, Finck R, Bruggner RV, Melamed R, Trejo A, Ornatsky OI, Balderas RS, Plevritis SK, Sachs K, Pe’er D, Tanner SD, Nolan GP. Single-Cell Mass Cytometry of Differential Immune and Drug Responses Across a Human Hematopoietic Continuum. Science. American Association for the Advancement of Science; 2011 May 6;332(6030):687–696.

32. Van Gassen S, Callebaut B, Van Helden MJ, Lambrecht BN, Demeester P, Dhaene T, Saeys Y. FlowSOM: Using self-organizing maps for visualization and interpretation of cytometry data. Cytometry Part A. 2015;87(7):636–645.

33. Nowicka M, Krieg C, Weber LM, Hartmann FJ, Guglietta S, Becher B, Levesque MP, Robinson MD. CyTOF workflow: differential discovery in high-throughput high-dimensional cytometry datasets. F1000Res. 2017 Nov 14;6:748.

34. McInnes L, Healy J, Melville J. UMAP: Uniform Manifold Approximation and Projection for Dimension Reduction [Internet]. arXiv; 2020 [cited 2025 Apr 21]. Available from: http://arxiv.org/abs/1802.03426

35. Zou H, Hastie T. Regularization and Variable Selection Via the Elastic Net. Journal of the Royal Statistical Society Series B: Statistical Methodology. 2005 Apr 1;67(2):301–320.

36. Kruskal WH, Wallis WA. Use of Ranks in One-Criterion Variance Analysis. Journal of the American Statistical Association. ASA Website; 1952 Dec;47(260):583–621.

37. Benjamini Y, Hochberg Y. Controlling the False Discovery Rate: A Practical and Powerful Approach to Multiple Testing. Journal of the Royal Statistical Society: Series B (Methodological). 1995;57(1):289–300.

38. Wilcoxon F. Individual Comparisons by Ranking Methods. In: Kotz S, Johnson NL, editors. Breakthroughs in Statistics: Methodology and Distribution [Internet]. New York, NY: Springer; 1992 [cited 2025 Feb 24]. p. 196–202. Available from: 10.1007/978-1-4612-4380-9_16

39. Guo N, Chen Y, Su B, Yang X, Zhang Q, Song T, Wu H, Liu C, Liu L, Zhang T. Alterations of CCR2 and CX3CR1 on Three Monocyte Subsets During HIV-1/Treponema pallidum Coinfection. Front Med [Internet]. Frontiers; 2020 Jun 18 [cited 2025 Mar 6];7. Available from: https://www.frontiersin.org/journals/medicine/articles/10.3389/fmed.2020.00272/full

40. Durda P, Raffield LM, Lange EM, Olson NC, Jenny NS, Cushman M, Deichgraeber P, Grarup N, Jonsson A, Hansen T, Mychaleckyj JC, Psaty BM, Reiner AP, Tracy RP, Lange LA. Circulating Soluble CD163, Associations With Cardiovascular Outcomes and Mortality, and Identification of Genetic Variants in Older Individuals: The Cardiovascular Health Study. J Am Heart Assoc. 2022 Oct 31;11(21):e024374. PMCID: PMC9673628

41. Aristoteli LP, Møller HJ, Bailey B, Moestrup SK, Kritharides L. The monocytic lineage specific soluble CD163 is a plasma marker of coronary atherosclerosis. Atherosclerosis. 2006 Feb 1;184(2):342–347.

42. Moreno JA, Ortega-Gómez A, Delbosc S, Beaufort N, Sorbets E, Louedec L, Esposito-Farèse M, Tubach F, Nicoletti A, Steg PG, Michel JB, Feldman L, Meilhac O. In vitro and in vivo evidence for the role of elastase shedding of CD163 in human atherothrombosis. Eur Heart J. 2012 Jan;33(2):252–263. PMID: 21606088

43. Etzerodt A, Moestrup SK. CD163 and Inflammation: Biological, Diagnostic, and Therapeutic Aspects. Antioxid Redox Signal. 2013 Jun 10;18(17):2352–2363. PMCID: PMC3638564

44. Subauste CS, de Waal Malefyt R, Fuh F. Role of CD80 (B7.1) and CD86 (B7.2) in the Immune Response to an Intracellular Pathogen1. The Journal of Immunology. 1998 Feb 15;160(4):1831–1840.

45. Thomas G, Tacke R, Hedrick CC, Hanna RN. Nonclassical Patrolling Monocyte Function in the Vasculature. Arteriosclerosis, Thrombosis, and Vascular Biology. American Heart Association; 2015 Jun;35(6):1306–1316.

46. Tahir S, Steffens S. Non-classical monocytes in cardiovascular physiology and disease. American Journal of Physiology-Cell Physiology [Internet]. American Physiological Society Rockville, MD; 2021 Feb 16 [cited 2025 Apr 22]; Available from: https://journals.physiology.org/doi/10.1152/ajpcell.00326.2020

47. Yona S, Lin HH, Dri P, Davies JQ, Hayhoe RPG, Lewis SM, Heinsbroek SEM, Brown KA, Perretti M, Hamann J, Treacher DF, Gordon S, Stacey M. Ligation of the adhesion-GPCR EMR2 regulates human neutrophil function. The FASEB Journal. 2008;22(3):741–751.

48. Altara R, Mallat Z, Booz GW, Zouein FA. The CXCL10/CXCR3 Axis and Cardiac Inflammation: Implications for Immunotherapy to Treat Infectious and Noninfectious Diseases of the Heart. J Immunol Res. 2016;2016:4396368. PMCID: PMC5066021

49. Mohapatra A, Howard Z, Ernst JD. CCR2 recruits monocytes to the lung, while CX3CR1 modulates positioning of monocyte-derived CD11c pos cells in the lymph node during pulmonary tuberculosis. bioRxiv. 2025 Feb 8;2025.02.07.637199. PMCID: PMC11839135

50. Sbrana S, Campolo J, Clemente A, Bastiani L, Cecchettini A, Ceccherini E, Caselli C, Neglia D, Parodi O, Chiappino D, Smit JM, Scholte AJ, Pelosi G, Rocchiccioli S. Blood Monocyte Phenotype Fingerprint of Stable Coronary Artery Disease: A Cross-Sectional Substudy of SMARTool Clinical Trial. BioMed Research International. 2020;2020(1):8748934.

51. Patel VK, Williams H, Li SCH, Fletcher JP, Medbury HJ. Monocyte inflammatory profile is specific for individuals and associated with altered blood lipid levels. Atherosclerosis. Elsevier; 2017 Aug 1;263:15–23. PMID: 28570862

52. SahBandar IN, Ndhlovu LC, Saiki K, Kohorn LB, Peterson MM, D’Antoni ML, Shiramizu B, Shikuma CM, Chow DC. Relationship between Circulating Inflammatory Monocytes and Cardiovascular Disease Measures of Carotid Intimal Thickness. J Atheroscler Thromb. 2020 May 1;27(5):441–448. PMCID: PMC7242227

53. Padgett LE, Araujo DJ, Hedrick CC, Olingy CE. Functional crosstalk between T cells and monocytes in cancer and atherosclerosis. Journal of Leukocyte Biology. 2020;108(1):297– 308.

54. Kim KW, Ivanov S, Williams JW. Monocyte Recruitment, Specification, and Function in Atherosclerosis. Cells. 2020 Dec 24;10(1):15. PMCID: PMC7823291

55. Li B, Shaikh F, Zamzam A, Abdin R, Qadura M. Inflammatory Biomarkers to Predict Major Adverse Cardiovascular Events in Patients with Carotid Artery Stenosis. Medicina. Multidisciplinary Digital Publishing Institute; 2024 Jun;60(6):997.

56. Wiche Salinas TR, Zhang Y, Gosselin A, Rosario NF, El-Far M, Filali-Mouhim A, Routy JP, Chartrand-Lefebvre C, Landay AL, Durand M, Tremblay CL, Ancuta P. Alterations in Th17 Cells and Non-Classical Monocytes as a Signature of Subclinical Coronary Artery Atherosclerosis during ART-Treated HIV-1 Infection. Cells. 2024 Jan 15;13(2):157. PMCID: PMC10813976

